# Actionable spatial prostanoid barriers constrain BiTE-driven adoptive T cell immunity in intact human tumors

**DOI:** 10.64898/2026.03.26.713601

**Authors:** Bovannak S. Chap, Tania Santoro, Paris Kosti, David Barras, Noémie Fahr, Mathieu Desbuisson, Fabrizio Benedetti, Aspram Minasyan, Asia Andreoli, Eleonora Ghisoni, Flavia De Carlo, Khadidja Benkortbi, Apostolos Sarivalasis, Chahin Achtari, Delfyne Hastir, Sabina Berezowska, Karim Abdelhamid, Christine Sempoux, Jean Yannis Perentes, Patrice Mathevet, Jose Ramon Conejo-Garcia, George Coukos, Steven M. Dunn, Evripidis Lanitis, Denarda Dangaj Laniti

## Abstract

Adoptive cell therapy (ACT) in solid tumors is limited by tumor microenvironment (TME)-imposed resistance mechanisms that are inadequately addressed by conventional systems. We developed tissue-preserving patient-derived explants (PDEs) from lung and ovarian cancer to interrogate redirected T cell immunity in intact human tissue. Using mesothelin-targeting bispecific T cell engager (BiTE^®^, Amgen trademark)-secreting T cells, we observed antigen-dependent but heterogeneous responses across lesions. An integrated *ex vivo* response score stratified responder and non-responder TMEs, revealing that resistance associates with reduced antigen density, stromal dominance, and limited myeloid licensing rather than baseline lymphocyte abundance. Elevated prostaglandin E_2_ (PGE_2_) inversely correlated with BiTE-induced T cell activation, identifying the COX/PGE_2_ axis as a tissue-imposed constraint. COX inhibition amplified interferon-driven immune programs enhanced intratumoral CD8⁺ infiltration, and increased tumor-restricted apoptosis. Spatial transcriptomics localized these effects to tumor-proximal immune hubs in responders, whereas non-responders remained stromally insulated. These findings position PDEs as human-based new approach methodologies enabling combinatorial ACT pharmacodynamics and stratification.

**Statement of significance:** Patient-derived explants provide a human-based new approach methodology to interrogate adoptive immunotherapy pharmacodynamics within intact tumor microenvironments in NSCLC and HGSOC. We uncover a COX/PGE_2_-mediated tissue ceiling that limits BiTE-driven T cell function and demonstrate that COX inhibition reactivates tumor-proximal immune hubs to enhance intratumoral CD8 infiltration and tumor-restricted apoptosis, informing patient stratification and rational combinations.

## Introduction

Adoptive cell therapy (ACT) can achieve durable tumor control in solid cancers, most notably with tumor-infiltrating lymphocytes (TILs) in melanoma, yet sustained benefit across most solid tumor indications remains uncommon. This gap reflects dynamic, heterogeneous tumor microenvironments (TMEs) that can constrain T cell trafficking, productive effector-target engagement, and local persistence. As tumors evolve, they diversify into spatially distinct molecular and ecological states with variable treatment sensitivity, contributing to relapse^1^.

A major barrier to improving ACT is that widely used preclinical models fail to preserve the spatial organization of the TME that governs immune control *in vivo*. In solid tumors, immune behavior is dictated within structured cell-cell and cell-matrix neighborhoods, where stromal and myeloid circuits shape T cell accumulation, and exposure to suppressive cues. Tissue dissociation disrupts these interactions and collapses critical gradients. In ovarian cancer (OC), immune-excluded and immune-desert phenotypes are supported by suppressive myeloid populations and stromal barrier states involving vasculature, cancer-associated fibroblasts (CAFs) and extracellular matrix (ECM) remodeling, with spatial profiling implicating tumor-stroma interfaces and matrisome-linked programs in poor-outcome niches^2–4^. These considerations have motivated *ex vivo* 3D platforms that preserve tissue architecture and patient-specific heterogeneity to enable controlled interrogation of immune-tumor crosstalk^4, 5^.

Conventional 2D and simplified 3D models (monolayers, spheroids, organoids) support mechanistic studies but do not capture patient-specific stromal and myeloid contexts or the ECM programs that shape clinical resistance^6, 7^. Patient-derived explants (PDEs) preserve native tumor architecture and multicellular TMEs, enabling short-term, spatially resolved pharmacodynamic readouts in intact human tissue^8, 9^. Despite this, PDEs have been applied only sparingly to evaluate engineered cellular therapies and microenvironment-targeted combinations^4^.

Among immunosuppressive pathways, the COX/PGE_2_ axis is a therapeutically tractable regulator of antitumor immunity that is associated with tumor progression and poor prognosis in NSCLC and HGSOC^10–13^. COX-derived PGE_2_ acts as a convergent mediator that limits both T cell fitness and myeloid activation^14, 15^. In human tumor-infiltrating lymphocytes (TILs) PGE_2_ signaling disrupts metabolic adaptation and mitochondrial resilience, curtailing effector T cell expansion and persistence during ACT^16, 17^. In parallel, PGE_2_ conditions the myeloid compartment toward tolerogenic dendritic cell states and protumorigenic macrophage polarization^12^. Together, these findings position the COX/PGE_2_ axis as an integrative suppressive node linking T cell-intrinsic dysfunction and myeloid-mediated immune exclusion^18^.

Aligned with increasing emphasis on human-based research and new approach methodologies (NAMs), we established tissue-preserving patient-derived explants (PDEs) as a human pharmacodynamic platform to functionally test adoptive immunotherapies within intact tumor microenvironments using PDEs from NSCLC and HGSOC. We focused on T cells engineered to secrete mesothelin (MSLN)-targeting bispecific T cell engagers (BiTEs), a diffusible CD3-engaging modality that redirects polyclonal T cells toward antigen-expressing targets and can recruit bystander T cells^19^. We hypothesized that BiTE-secreting T cells initiate antigen-dependent activation within tumor islets but that efficacy is limited by prostanoid-associated stromal and myeloid barrier states. In responsive lesions, MSLN-BiTE T cell engagement induces interferon, chemokine, and cytotoxic programs and that enable tumor-restricted killing, whereas non-responsive lesions retain COX/PGE_2_-linked stromal and vascular programs that prevent immune-hub formation. COX inhibition is predicted to relieve this constraint and amplify engineered and endogenous T cell responses.

Using tiered 3D tumor models and TME-preserving PDEs, we defined how antigen abundance and tissue architecture shape BiTE-T cell cytotoxicity, tested whether COX/PGE_2_ signaling caps early *ex vivo* responses, and resolved spatial immune activation versus barrier programs that distinguish responder and non-responder lesions. Together, these data link intact human tissue pharmacodynamics with a defined ACT modality and a targetable suppressive pathway to inform response prediction and rational combination design in solid tumors.

## Results

### Patient-derived tumoroids reveal antigen-dependent potency of adoptive T cell therapies in 3D tumors

To assess engineered tumor-reactive T cells in architectures that capture diffusion and geometric constraints, we established 3D models ^20^ instead of monolayer co-cultures. We first developed a tiered system using OVCAR-8 monolayers and Matrigel-free OVCAR-8 spheroids, and compared two MHC-independent MSLN-targeted products ^19^, a second-generation MSLN-CAR and MSLN-BiTE T cells (**Fig. 1A, Fig. S1A**) having relatively comparable transduction efficiencies and being dominated by CD8 T cells (**Fig. S1B-C)**. Effector and target cells were co-cultured for 24 hrs at an estimated 1:1 effector: target (E:T) ratio, and reactivity was quantified by CD137⁺ (4-1BB) upregulation^21^, and IFNγ, TNFα, and granzyme B secretion.

**Figure 1:**
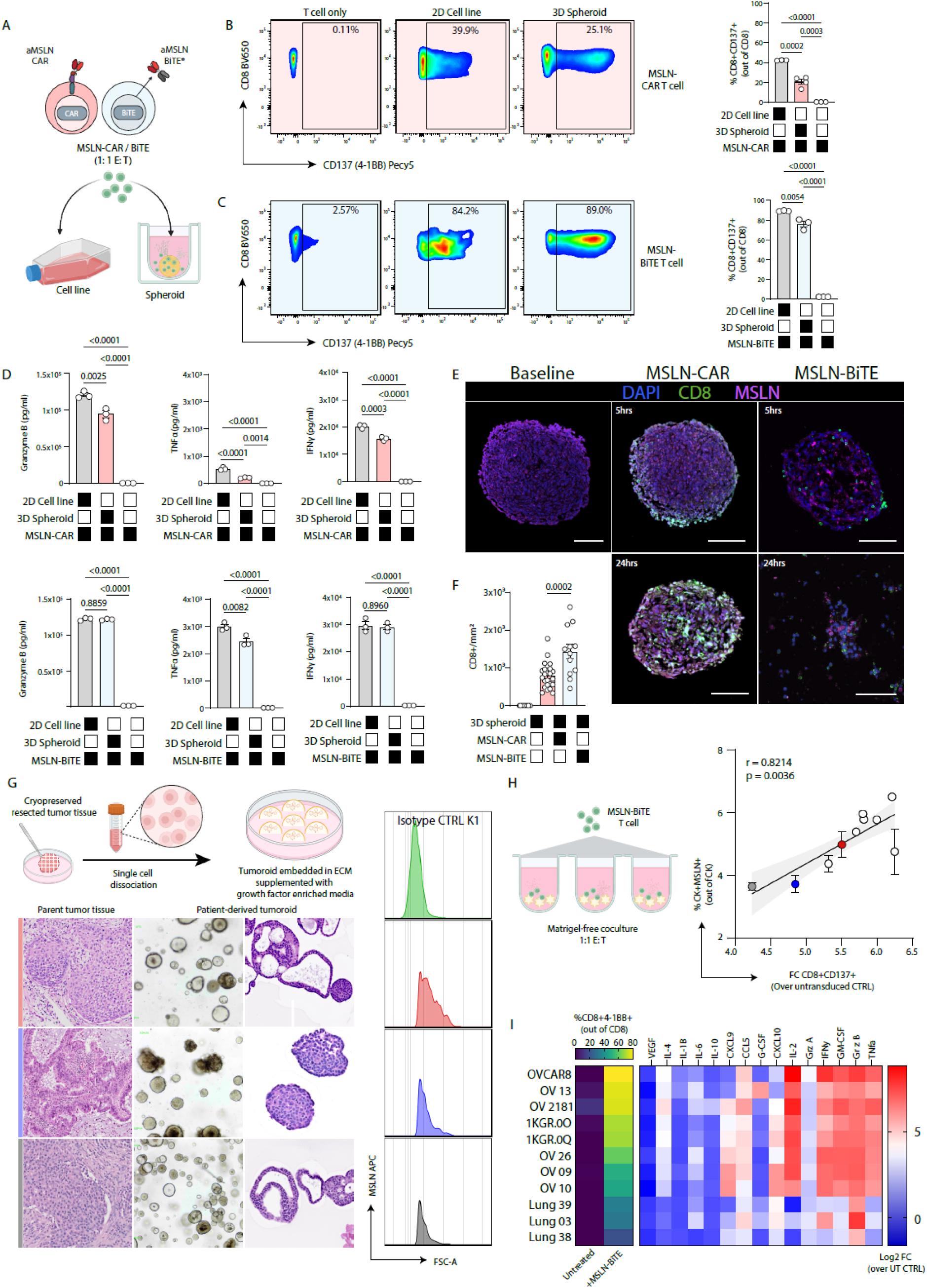
Patient-derived spheroids and tumoroids enable functional benchmarking of MSLN-targeted adoptive T cell products in 3D. **A,** Schematic of the tiered benchmarking assay comparing anti-MSLN CAR-T cells and anti-MSLN BiTE T cells in OVCAR-8 2D monolayers and matrix-free 3D spheroids after 24 hrs coculture at a 1:1 effector-to-target (E:T) ratio. **B,** Representative flow cytometry plots and quantification of CD8^+^CD137^+^ cells among CD8^+^ T cells after coculture of MSLN-CAR T cells alone (T cell only), with 2D OVCAR-8 cells, or with 3D OVCAR-8 spheroids. **C,** Representative flow cytometry plots and quantification of CD8^+^CD137^+^ cells among CD8+ T cells after coculture of MSLN-BiTE T cells alone (T cell only), with 2D OVCAR-8 cells, or with 3D OVCAR-8 spheroids. **D,** Supernatant concentrations of granzyme B, TNFα, and IFNγ after 24-hour coculture of MSLN-CAR (top) or MSLN-BiTE (bottom) T cells with 2D OVCAR-8 cells or 3D spheroids. **E,** Representative confocal immunofluorescence images of embedded 3D spheroids stained for nuclei (DAPI), CD8, and MSLN at baseline and at 5 and 24 hrs after exposure to MSLN-CAR or MSLN-BiTE T cells. Scale bars, 100 μm. **F,** Quantification of intratumoral CD8 density in 3D spheroids across conditions. Statistical analysis using one-way ANOVA (+Kruskal-Wallis) **G,** Quantification of intratumoral CD8 density in 3D spheroids across conditions. **H,** Interpatient heterogeneity of MSLN expression in PDTs assessed by flow cytometry, schematic of Matrigel-free PDT coculture with MSLN-BiTE T cells at a 1:1 E:T ratio, and correlation between epithelial target abundance (%CK^+^MSLN^+^ of CK^+^ cells) and CD8 activation (FC of CD8^+^CD137^+^ over UT control). **I**, Multiparametric functional profiling of PDT responses to MSLN-BiTE T cells. Upper part, CD8 activation (%CD8⁺CD137⁺) under untreated versus +MSLN-BiTE conditions. Below, heatmap summarizing changes in secreted mediators (log2 fold-change over UT control) following co-culture across the indicated tumoroid lines. One-way repeated-measures ANOVA (+Tukey) was performed in GraphPad Prism v10.2 (n=3-4 technical replicates per patients). Exact P values and comparisons are indicated in the panels. In **B** and **C**, representative plots are shown and quantification reflects 3 to 4 technical replicates per condition. In **D**, *n* = 3 technical replicates per condition. In **F**, *n* = 12 to 22 regions of interest per condition. Exact *P* values and comparisons are indicated in the panels. Figure created with BioRender.

MSLN CAR-T cells displayed consistently lower reactivity when cocultured in a 3D spheroid setting compared to the 2D monolayer coculture (**Fig. 1B-C**). As previously shown MSLN-BiTE T cells outperformed MSLN-CAR T cells against OVCAR-8 cells cultured in 2D monolayers^19^. MSLN-BiTE consistently elicited higher expression of CD137⁺ (4-1BB) in CD8⁺ T cells (**Fig. 1B**), and a greater release of IFNγ, TNFα, and granzyme B than MSLN-CAR T cells in the 2D cocultures (**Fig. 1D, light blue**).

Notably, this superior performance of MSLN-BiTE T cells persisted in OVCAR-8 spheroid cocultures, where MSLN-BiTEs maintained robust activation (∼80%) and inflammatory mediators secretion comparable to the monolayer cocultures, whereas CAR T cells exhibited attenuated CD137^+^ up-regulation (∼20%) with modest reductions in IFNγ, TNFα, and granzyme B relative to the 2D system (**Fig. 1D, pink**). These first data suggested that in contrast to CAR-T cells, CD3ε-engaging BiTEs could better penetrate tumor interstices and potentially have improved infiltration in 3D tumors.

Therefore, we sought to monitor the infiltration capacity of both MSLN-CAR and MSLN-BiTE T cells in 3D OVCAR8 spheroids after longitudinal cocultures. We embedded our samples from our 3D cocultures in paraffin and performed confocal imaging on serial sections of the 3D embedded spheroids (**Fig. 1E, confocal imaging IF**). In the MSLN-BiTE condition, CD8⁺ T cells progressively infiltrated and disrupted the spheroids, culminating in a complete eradication of the 3D tumor at 24hrs (**Fig. 1E**). In contrast, quantification of CAR T cells showed lower intratumoral distribution and infiltration after 5hrs coculture (**Fig. 1F**). This data indicates that MSLN-BiTEs not only sustain effector activation in 3D tumors but also promote more effective tissue penetration.

While OVCAR-8 spheroids allow us to dissect biomechanical constraints, they represent clonotypic and homogeneous structures of cancer cells. Indeed, the target antigen MSLN was highly and equivalently expressed in 2D and 3D cell culture formats (**Fig. S1D).** To interrogate MSLN-BiTE functionality in more clinically relevant settings, we established patient-derived tumoroids (PDTs) from NSCLC and HGSOC, which are expected to preserve epithelial architecture^22–25^ while potentially recapitulating heterogeneous MSLN expression across individuals. Tumoroids from lung and ovarian tissues were generated using Matrigel and growth factor-supplemented media (**Fig. 1G & methods**). Hematoxylin and eosin (H&E) staining and brightfield imaging confirmed epithelial morphology, while flow cytometry analysis revealed inter-patient heterogeneity in MSLN expression (**Fig. 1G, histograms**).

Upon establishment and propagation, our PDTs were transitioned in Matrigel-free 3D cultures to enable coculture with T cells in a matrix-free environment, alike the OVCAR8 spheroid system. We then cocultured NSCLC and HGSOC PDTs with MSLN-BiTE and control untransduced (UT) T cells for 24hrs, and brightfield imaging revealed patient-dependent differences in tumoroid disruption in the MSLN-BiTE condition (**Fig. S1E**). MSLN-BiTE T cells were reactive against PDTs and we found that their activation, as measured by CD137⁺ expression, was highly correlated with the levels of MSLN expression by the tumoroids (**Fig. 1H**). Secretome analysis at 24 hrs confirmed this cellular readout: granzyme B, IFNγ, and TNFα were consistently elevated in 3D PDT co-cultured with MSLN-BiTEs (**Fig. 1I**).

These results indicate that 3D epithelial models, such as spheroids and PDTs, provide sensitive, quantitative readouts of redirected T cell function. Nevertheless, PDTs lack the full cellular and biomechanical complexities of the TME, particularly stromal and myeloid populations that critically modulate T cell recruitment and their effector durability^26, 27^. Consistent with this limitation, the secretome of PDTs were mainly dominated by T cell-derived cytokines (IFNγ, TNFα, IL-2) with little representation of stromal/myeloid cytokines or chemokines (**Fig. 1H**). For example, CXCL9/10 levels remained near baseline levels despite robust IFNγ and TNFα induction (**Fig. 1H**).

Together, these findings establish patient-derived tumoroids as sensitive 3D platforms to functionally benchmark adoptive T cell products, capturing antigen dependence, tissue penetration, and cytotoxic efficacy across solid tumor types. However, the limited stromal and myeloid representation in tumoroids underscores the need for complementary TME-preserving models to fully predict immunotherapy performance in patients.

### Patient-derived explants faithfully preserve native tumor-immune architecture *ex vivo*

A body of evidence indicating that intratumoral myeloid cells from the TME orchestrate the ACT response prompted us to investigate the MSLN-BiTE product within a more complex preclinical model that maintains epithelial, stromal and immune compartments^12, 26, 28^. Therefore, to assess the immunological response of the MSLN-BiTEs in a patient-relevant preclinical model, we set up an organotypic coculture platform using patient-derived explants (PDE) which permit *ex vivo* perturbations that could remodel local immunity^4, 5^.

We focused our study on NSCLC and HGSOC PDEs because both diseases express MSLN in a clinically meaningful fraction. HGSOC shows frequent positivity (̃55-97%), while NSCLC, particularly adenocarcinoma, harbors MSLN in a substantial subset (̃24-69%)^29–31^.

Sixty-nine samples from HGSOC and NSCLC specimens were obtained by surgical resection, and cryopreserved in our biobank. To move forward we introduced a standardized qualification pipeline prior to their inclusion in downstream PDE experiments (**Fig. 2A**).

**Figure 2:**
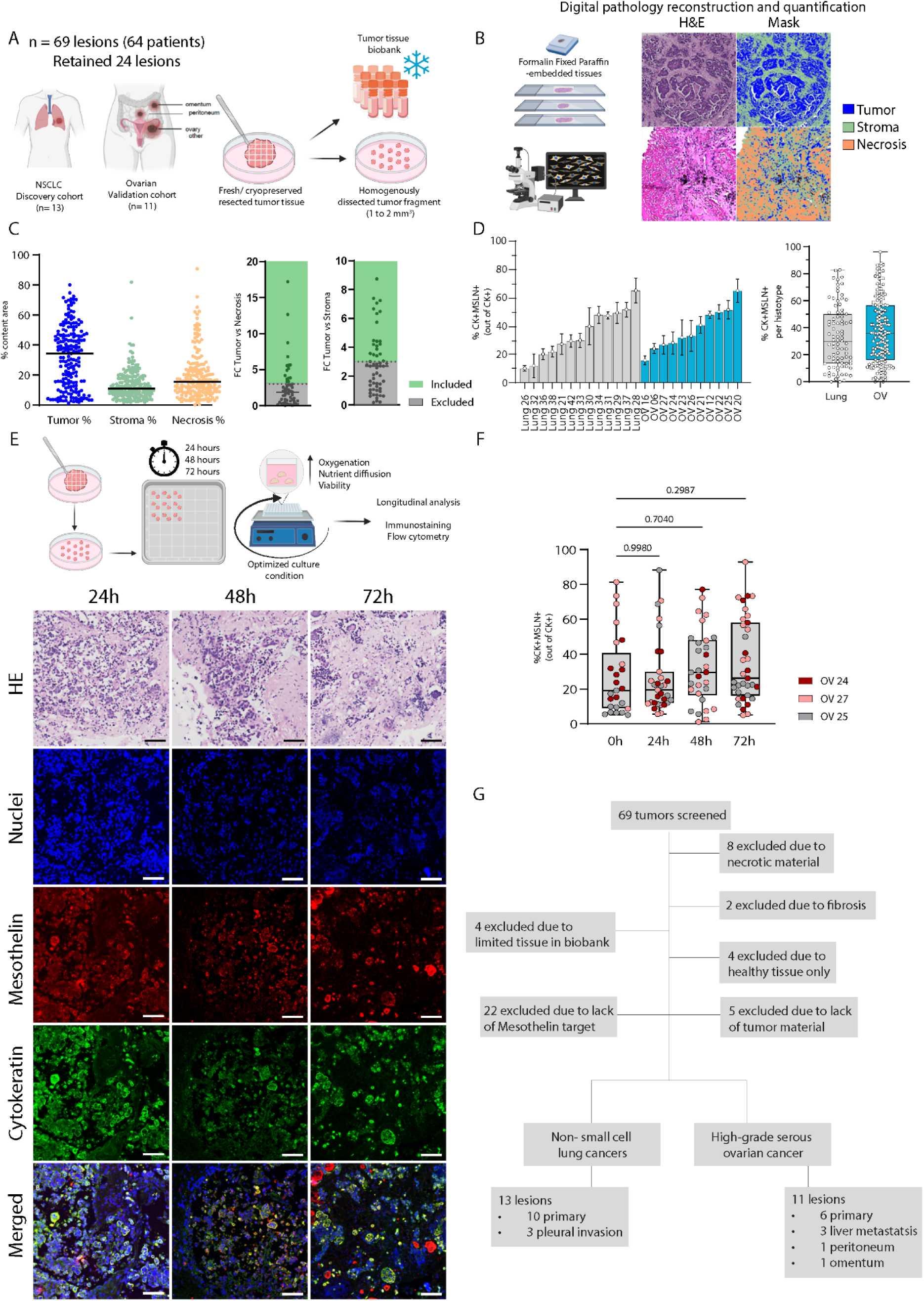
Standardized qualification pipeline for NSCLC and HGSOC PDEs to preserve tumor architecture and MSLN expression within a defined functional window. **A,** Overview of the histoculture workflow and cohort composition. Surgically resected NSCLC and HGSOC specimens were collected fresh or from tumor biobank cryostocks and processed into uniformly dissected tumor fragments of approximately 1 to 2 mm³ for downstream *ex vivo* culture. A total of 69 lesions from 64 patients were screened, and 24 lesions were retained for functional PDE experiments. **B,** Digital pathology reconstruction and quantitative tissue segmentation workflow. Formalin-fixed, paraffin-embedded (FFPE) sections were stained with H&E and computationally segmented into tumor, stroma, and necrosis compartments to generate lesion-level tissue composition metrics. **C,** Distribution of tumor, stroma, and necrosis fractions across screened lesions based on digital pathology quantification (**left**). **Right**, inclusion criteria based on relative tissue composition, shown as fold change of tumor area over necrosis and stroma, identifying lesions retained or excluded according to predefined thresholds. **D,** Baseline MSLN target abundance in retained lesions, quantified by IF as the percentage of CK^+^MSLN^+^ cells among CK^+^ epithelial cells. **Left**, lesion-level values across individual NSCLC and HGSOC samples. **Right**, pooled values by histotype. **E,** Determination of the PDE culture window. Top, schematic of longitudinal assessment performed at 24, 48, and 72 hrs under optimized culture conditions. Bottom, representative H&E and immunofluorescence images acquired at each time point for nuclei (DAPI), MSLN, CK, and merged channels, showing preservation of epithelial architecture and target expression over the analyzed interval. **F,** Longitudinal quantification of MSLN expression stability during culture, shown as the percentage of CK^+^MSLN^+^ cells among CK^+^ epithelial cells at 0, 24, 48, and 72 hrs in three independent lesions. Dots represent regions of interest. **G,** Diagram summarizing screening and exclusion steps applied to the 69 screened lesions, including exclusions due to insufficient tumor material, excessive necrosis or fibrosis, healthy tissue only, and lack of mesothelin expression, resulting in the final set of NSCLC and HGSOC PDEs used for downstream functional assays. In **F**, statistics were performed on patient-level means using one-way repeated-measures ANOVA with Dunnett multiple-comparisons test versus 0 hr. Exact P values are indicated in the panel. Figure created with BioRender.

First, we performed multiparameter flow cytometry of our 69 samples to profile TME composition at the single cell level. We quantified the frequencies of EPCAM⁺ epithelial tumor cells (including the EPCAM⁺MSLN⁺ compartment), EPCAM⁻CD45⁻ stromal or other cells, as well as CD45⁺ immune populations comprising CD3⁺ T cells (CD4⁺ and CD8⁺ subsets and CD137⁺ activated fractions), CD19⁺ B cells, NK cells, and myeloid subsets including CD14⁺ cells, CD68⁺ macrophages (with CD80⁺ and CD206⁺), and CD11c⁺ dendritic cells (**Fig. 2SA-C**).

We observed that epithelial cancer cell fractions measured by EpCAM expression varied widely with a substantial fraction of specimens in both HGSOC and NSCLC cohorts containing mostly stromal and immune cells. Those cases were excluded from further use as PDEs.

Then, to ensure architectural and cellular adequacy for downstream assays, we implemented quantitative digital pathology on H&E and IF section to segment tissues of each PDE into tumor, stroma, and necrotic areas (**Fig. 2B-C**) and quantify MSLN expression (**Fig. 2D, S2D**). These metrics were then used to screen PDEs for functional cocultures (**Methods**). Our PDE model inclusion criteria required > 3-fold higher tumor area relative to stroma and necrosis and detectable MSLN expression by IF (**Fig. 2C-D**).

We next defined the culture window during which our histocultures (1-2 mm³) preserve their original structure. Using three independent patients (multiple fragments; n= 6 lesions per timepoint and per patient to capture intra-patient heterogeneity), we profiled PDEs after 24, 48, and 72 hrs by immunofluorescence (IF) and flow cytometry (**Fig. 2E-F, S2A-D**). We quantified antigen expression in differentiated epithelial cells (defined as the percentage of MSLN⁺ among CK⁺ cells) and found that they remained stable over time (**Fig. 2F).** To assess potential changes in immune cellular composition within the PDE, we performed flow cytometry on dissociated explants to quantify leukocyte subsets (CD3⁺, NK cells, B cells and myeloid populations) longitudinally (**Fig. S2E**).

Across samples, myeloid compartments (CD11c⁺ and CD68⁺ cells) remain largely stable, whereas the fractions of lymphoid populations were very dynamic over time (**Fig. S2E**). To minimize culture-induced drift and best preserve the TME cellular composition we therefore fixed the operative timeframe for functional experiments to 24 hrs, as this timepoint shows minimal deviation compared with later culture timepoints. Of the 69 lesions from 64 patients initially screened, 24 specimens (NSCLC = 13, HGSOC = 11) passed our inclusion criteria (**Fig. 2G** and **Methods**).

Together, these data establish PDEs as a robust and stringently qualified 3D platform that maintains tumor architecture, mesothelin expression, and key immune and stromal compartments within a defined experimental window. This system enables controlled, patient-relevant interrogation of redirected T cell activity within an intact and biologically authentic TME.

### Mesothelin-dependent BiTE activity triggers a coordinated IFNγ-chemokine-cytotoxic response in PDEs

To directly investigate whether MSLN antigen expression in PDEs can be leveraged to drive early, MSLN-BiTE T cell activation *in situ,* PDEs with MSLN expression were employed for functional T cell coculture assays. Co-cultures included three conditions: untreated PDEs (medium only), PDE cocultured with UT T cells (negative control) and PDE cocultured with MSLN-BiTE T cells (**Fig. 3A** and **Methods**).

**Figure 3:**
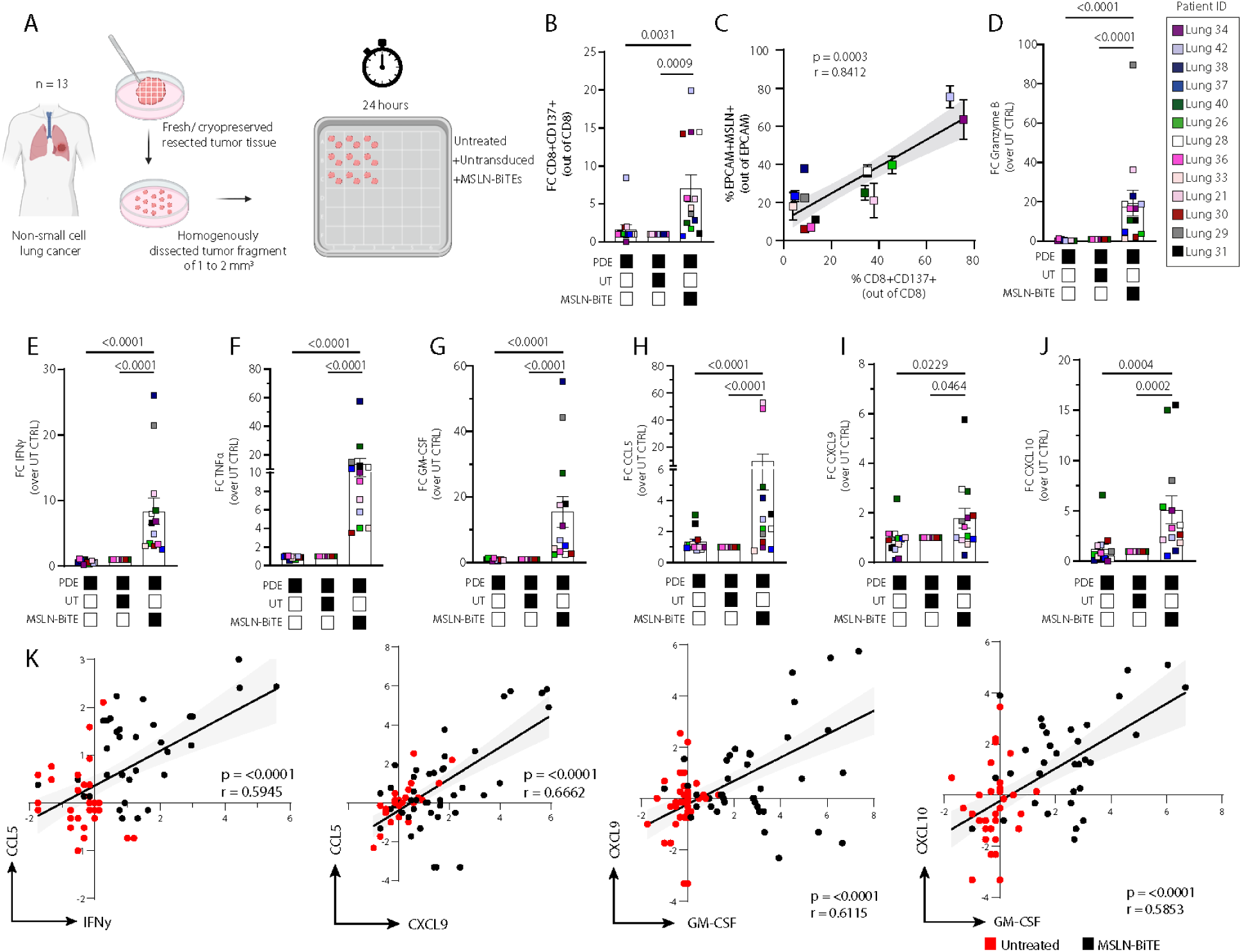
MSLN-BiTE induces tumor-reactive CD8⁺ activation and a coordinated inflammatory/chemokine secretome in NSCLC PDEs. **A,** Schematic of the NSCLC patient-derived explant (PDE) assay. Fresh or cryopreserved resected tumor tissue from 13 patients was dissected into homogeneous 1 to 2 mm³ fragments and cultured for 24 hours under untreated, untransduced (UT), or MSLN-BiTE conditions. **B,** FC in the frequency of activated CD8⁺CD137⁺ T cells (out of total CD8⁺) across conditions (P values shown). **C,** Correlation between epithelial target abundance, quantified as the percentage of EpCAM^+^MSLN^+^ tumor cells among EpCAM^+^ cells, and ex vivo T-cell activation, quantified as the frequency of CD8^+^CD137^+^ cells among CD8^+^ T cells. **D,** FC of granzyme B relative to the UT control condition. **E-J,** FC of soluble mediators relative to the UT control condition: IFNγ (**E**), TNFα (**F**), GM-CSF (**G**), CCL5 (**H**), CXCL9 (**I**), and CXCL10 (**J**). symbols are color-coded by lesion/patient ID; bars indicate mean ± SEM. Statistical testing was performed on lesion-level values using one-way repeated-measures ANOVA with appropriate multiple-comparisons correction (as indicated by the plotted comparisons). Exact P values are displayed on the graphs (GraphPad Prism v10.1.2). **K,** Pairwise correlation analyses linking key components of the inflammatory and chemokine response program, including CCL5 versus IFNγ, CCL5 versus CXCL9, CXCL9 versus GM-CSF, and CXCL10 versus GM-CSF. Red and black points denote untreated and MSLN-BiTE conditions, respectively. In **B-J**, points represent individual patients color-coded; bars indicate mean ± SEM. Exact *P* values are shown in the panels. Statistical testing was performed at patient-level values using one-way repeated-measures ANOVA with multiple-comparisons correction, as indicated by the plotted comparisons. In **K**, each point represents one explant fragment.

Interestingly, flow cytometric analysis of NSCLC PDEs cultured in media only showed basal levels of CD137 expression in endogenous TIL within the PDE (**Fig. S3A**). Analysis of NSCLC PDEs cocultures revealed a significant increase in the frequency of intratumoral CD8⁺CD137^+^ T cells exclusively in the MSLN-BiTE condition compared to the untreated or UT T cell coculture control conditions (**Fig. 3B, Fig. S3A**). Across patients and PDEs, higher intratumoral CD8⁺CD137^+^ T cell frequencies strongly correlated with MSLN⁺ expression in CK⁺ NSCLC cells (**Fig. 3C, Fig. S3B**), indicating an antigen-dependent activation of MSLN-BiTE T cells.

Quantitative analysis of supernatants 24 hrs post PDE co-culture with MSLN-BiTE T cells showed robust induction of effector molecule secretion (granzyme B, IFNγ, TNFα, CCL5, CXCL9/10) relative to untreated PDEs or PDEs co-cultured with UT T cells in about 50% of our NSCLC patient cohort (**Fig. 3D-J, Fig.S3C-G**). Furthermore, correlation analyses of cytokines and chemokines secretion revealed highly coordinated upregulation of immune mediators: CCL5 positively correlated with IFNγ (Pearson r = 0.595, p < 0.0001) and CXCL9 (r = 0.666, p < 0.0001), while both CXCL9 and CXCL10 significantly correlated with GM-CSF (r = 0.612 and r = 0.586, respectively; p < 0.0001 for both) (**Fig. 3K**).

Together, these findings demonstrate that MSLN expression within PDEs is a key determinant of BiTE-mediated T cell activation, driving antigen-dependent expansion of intratumoral CD8⁺CD137⁺ effector cells. The tightly correlated induction of IFNγ, chemokines, and cytotoxic mediators reveals a coordinated immune program consistent with effective *in situ* engagement and amplification of anti-tumor immunity within the native TME.

### COX/PGE_2_ blockade rewires the tumor microenvironment to unleash BiTE-driven T cell immunity

We observed that in some PDEs despite expressing MSLN, MSLN-BiTE T cells failed to elicit anti-tumor responses (**Fig. S3B**). We hypothesized that TME-intrinsic suppressive mediators could limit MSLN-BiTE T cell reactivity. We previously demonstrated that prostaglandin E2 (PGE_2_), can dampen T cell effector function^17^. We therefore questioned if PGE_2_ was readily produced in our PDEs and quantified its secretion from each explant. Overall, the PGE_2_ concentrations fell in previously reported ranges (1nM to 1 µM) with a marked variation between patients (**Fig. 4A, heatmap**). Interestingly we found that the higher the PGE_2_ levels were in our PDE models the fewer were the frequencies of activated CD8⁺CD137⁺ T cells infiltrating them (**Fig. 4B**). PGE_2_ levels also inversely correlated with granzyme B release post MSLN-BiTE T cell cocultures (**Fig. 4C**), indicating that PGE_2_-rich milieus blunt synthetic T cell activation^17^.

**Figure 4:**
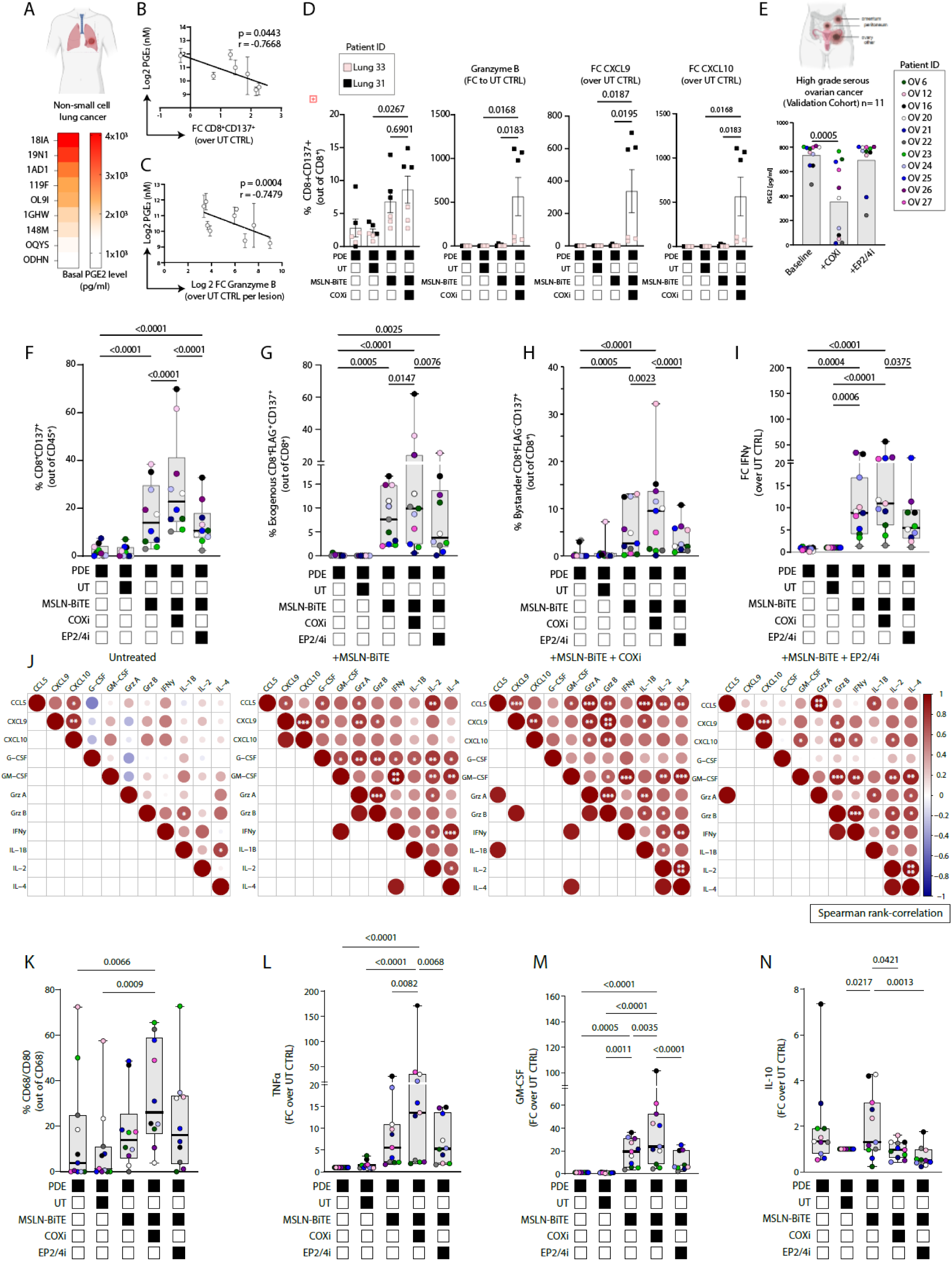
COX inhibition relieves a PGE_2_-imposed ceiling on MSLN-BiTE T cell activity and promotes inflammatory remodeling in NSCLC and HGSOC PDEs. **A,** Basal PGE_2_ quantified in NSCLC PDE supernatants (pg/mL), shown as a heatmap illustrating inter-lesion variability. **B,** Correlation between basal PGE_2_ and MSLN-BiTE-induced CD8+ T cell activation in NSCLC PDEs, expressed as FC in CD8^+^CD137^+^ frequencies relative to UT CTRL (Holm-corrected P value shown). **C,** Correlation between basal PGE_2_ and MSLN-BiTE-induced granzyme B release in NSCLC PDEs, expressed as log2 FC relative to UT CTRL per patient (r and P value shown). **D,** COX blockade rescue experiment in two NSCLC cases (Lung 31 and Lung 33): PDEs were pretreated with ketorolac (COXi, 1 µM) for 1 h prior to co-culture with MSLN-BiTE T cells, and CD8^+^CD137^+^ activation together with granzyme B, CXCL9, and CXCL10 were quantified as FC over UT CTRL. **E,** Pharmacodynamic verification of prostanoid pathway inhibition in the HGSOC validation cohort (n = 11), showing PGE_2_ levels at baseline and after COXi (ketorolac, 1 µM) or EP2/EP4 inhibition (EP2/4i, 1 µM). **F,** CD8^+^ T cell activation in HGSOC PDE co-cultures, reported as %CD8^+^CD137^+^ among CD8^+^ cells across conditions (condition matrix shown). **G-H,** FLAG-based discrimination of engineered versus bystander/resident CD8^+^ T cells in HGSOC PDEs, showing frequencies of CD8^+^CD137^+^FLAG^+^ *exogenous* cells (**G**) and CD8^+^CD137^+^FLAG^-^bystander/*endogenous* cells (**H**). (I) IFNγ secretion in HGSOC PDE co-cultures, expressed as FC over UT CTRL across conditions. (**J**) Cytokine and chemokine coordination across HGSOC PDEs visualized as Pearson correlation matrices for untreated, MSLN-BiTE, MSLN-BiTE + COXi, and MSLN-BiTE + EP2/4i conditions (circle size indicates correlation magnitude; color indicates direction). **K,** Myeloid polarization in HGSOC PDEs quantified as %CD68^+^CD80^+^ among CD68^+^ macrophages across conditions. (**L-N**) TNFα (**L**), GM-CSF (**M**), and IL-10 (**N**) secretion in HGSOC PDE co-cultures, expressed as FC over UT CTRL. Boxplots show median and interquartile range with whiskers as displayed; each dot represents one patient. For HGSOC panels (**E-N**), statistics were performed on patient-level values using one-way repeated-measures ANOVA with multiple-comparisons correction as indicated (GraphPad Prism v10.1.2). Figure generated with BioRender.

We decided to use this NSCLC discovery cohort as a test to determine if blockade of PGE_2_ synthesis could restore antitumoral activity. To block PGE_2_ production by the TMEs, we preconditioned PDEs from 2 patients (**lung 31 & 33**) with 1 µM of cyclooxygenase inhibitor (COXi = Ketorolac) 2 hrs prior to the addition of MSLN-BiTE T cells. Although there was a high variation between PDEs of the 2 patients included, we still observed a drastic increase in granzyme B secretion post COX inhibition and therefore a restoration of MSLN-BiTE T cell activity (**Fig. 4D)**. This response was accompanied by a 20-60 fold increase in CXCL9/10 production by the PDEs cocultured with MSLN-BiTE T cells in the presence of COXi (**Fig. 4D, lung 31**). Our data strongly suggest that COX inhibition *in situ* can relieve prostanoid-mediated suppression of immune cells in PDEs therefore potentiating the anti-tumor efficacy of engineered BiTE T cells.

To validate our findings, we employed PDEs from our HGSOC cohort. This time, in addition to COXi we added in our setting 1 µM of a selective EP2/EP4 inhibitor before coculture with MSLN-BiTE T cells, to block PGE_2_ receptors in T cells. To verify blockade of COX enzymes in the same PDEs used for functional readouts, we quantified PGE_2_ in supernatants after cocultures. COXi consistently reduced PGE_2_ concentration for most PDEs, lowering the cohort’s median by more than 50% relative to baseline (**Fig. 4E)**. As expected, EP2/EP4 inhibition had no effect on PGE synthesis.

When coculturing our HGSOC PDEs with irrelevant or MSLN-BiTE T cells we obtained equivalent infiltration levels between UT or MSLN-BiTE T cells in the PDEs, suggesting that the motility of exogenous T cells was not impacted by the TME (**Fig. S4A**). Also across cocultures of PDEs from HGSOC with MSLN-BiTEs, we observed that the total fraction of CD8⁺ cells upon inhibition of the COX/PGE_2_ was only slightly greater than without COXi (**Fig. S4A**). Importantly, and alike in the NSCLC setting, the reactivity of MSLN-BiTE T cells against PDEs, as measured by CD137⁺ expression, was significantly potentiated by COXi but not by inhibition of EP2/4 receptors (**Fig. 4F**).

We then asked whether exogenous MSLN BiTE-secreting T cells could mobilize and trigger the activation of endogenous tumor-resident T cells within the PDEs. To distinguish engineered BiTE-secreting T cells from endogenous tumor-infiltrating T lymphocytes (TILs) within PDEs, we took advantage of a FLAG-tag epitope incorporated into the MSLN-BiTE construct (**Fig. S4B**). FLAG⁺ marks transduced MSLN BiTE-secreting T cells, whereas FLAG⁻ identifies bystander and PDE-resident TILs.

We then quantified the proportions of exogenous MSLN-BiTE CD8⁺CD137⁺FLAG^+^ and bystander CD8⁺CD137⁺FLAG^-^ T cells. Relative to UT or PDE alone conditions, both the addition of MSLN-BiTEs T cells and their combination with COXi, increased the proportions of endogenous and exogenous CD8⁺CD137⁺ compartments (**Fig. 4G-H, Fig. S4C-D**). Effector function captured by IFNγ, TNFa, GM-CSF, CCL5, IL-2 and other cytokines was highly potentiated by COXi (**Fig. 4I and Fig. S4E-J**) at patient (**Fig. 4I-J**) and PDE level (**Fig. S4E-J**).

To infer coordination among soluble mediators, we computed pairwise Spearman rank-correlation across all HGSOC PDES for all measured cytokines and chemokines (**Fig. 4J**). Cocultures with MSLN-BiTE T cells assembled a Th1-centered module where IFNγ, CXCL9, CXCL10, and CCL5 covaried and linked to granzyme B reflecting a T cell centric modulation. Addition of COX inhibition strengthened and broadened this architecture, integrating GM-CSF and IL-1β into the same positively correlated cluster, a pattern indicating myeloid cell activation. Of note, EP2/EP4 antagonism increased the IFNγ/CXCR3-ligand node but yielded a less integrated network than COXi to the MSLN-BiTE cocultures (**Fig. 4J**).

We then asked whether this soluble signature coincided with phenotypic modulation of PDE-resident myeloid cells. COXi significantly increased CD68⁺CD80⁺ TAMs, when compared to MSLN-BiTE alone or EP2/EP4 inhibition (**Fig. 4K)**. This was supported by an increase of TNFα and GM-CSF secretion (**Fig. 4L-M**), along with a reduction of IL-10 (**Fig. 4N**) and VEGFA (**Fig. S4M**) immunoregulatory factors frequently associated with suppressive myeloid and regulatory immune programs in tumors. Together, these data demonstrate that upstream COX inhibition overcomes prostanoid-mediated immunosuppression, amplifying both engineered and endogenous T cell responses while reprogramming myeloid cells toward a pro-inflammatory state. By simultaneously reducing PGE_2_ availability and dampening its downstream signaling across multiple cellular compartments, COX blockade outperforms selective EP2/EP4 inhibition and emerges as a powerful strategy to potentiate BiTE-based adoptive T cell therapy.

### Spatial imaging unveils intratumoral CD8⁺ engagement and tumor-restricted killing in ovarian patient-derived explants

Building on the coordinated IFNγ-CXCL9/10 chemokine program and enhanced CD8⁺ activation observed upon COX inhibition (**Fig. 4J**), we asked whether these functional signatures translate into bona fide intratumoral recruitment and engagement of redirected T cells. High-parameter flow cytometry revealed increased CD8⁺ accumulation within PDEs following COX blockade. However, this approach could not resolve whether infiltrating cells were truly embedded within tumor regions or merely associated with the explant periphery. Although, UT T cells also exhibited increased apparent infiltration following explant dissociation and flow cytometric analysis (**Fig. 4G**), we could not exclude the possibility that these measurements reflected peripheral adhesion rather than true intratumoral engagement.

To distinguish true intratumoral localization from passive association, we performed volumetric two-photon microscopy following overnight coculture of PDEs with MSLN-BiTE T cells from two NSCLC and two HGSOC patients (30 lesions in total). Prior to coculture with PDEs, MSLN-BiTE T cells were fluorescently labeled to enable their tracking within the explants. Fixed whole-mount PDEs were stained for cytokeratin with a panCK antibody to define the tumor regions, DAPI for stroma, and subjected to tissue clearing before volumetric imaging by two-photon microscopy (**Fig. 5A, Msethods**).

**Figure 5:**
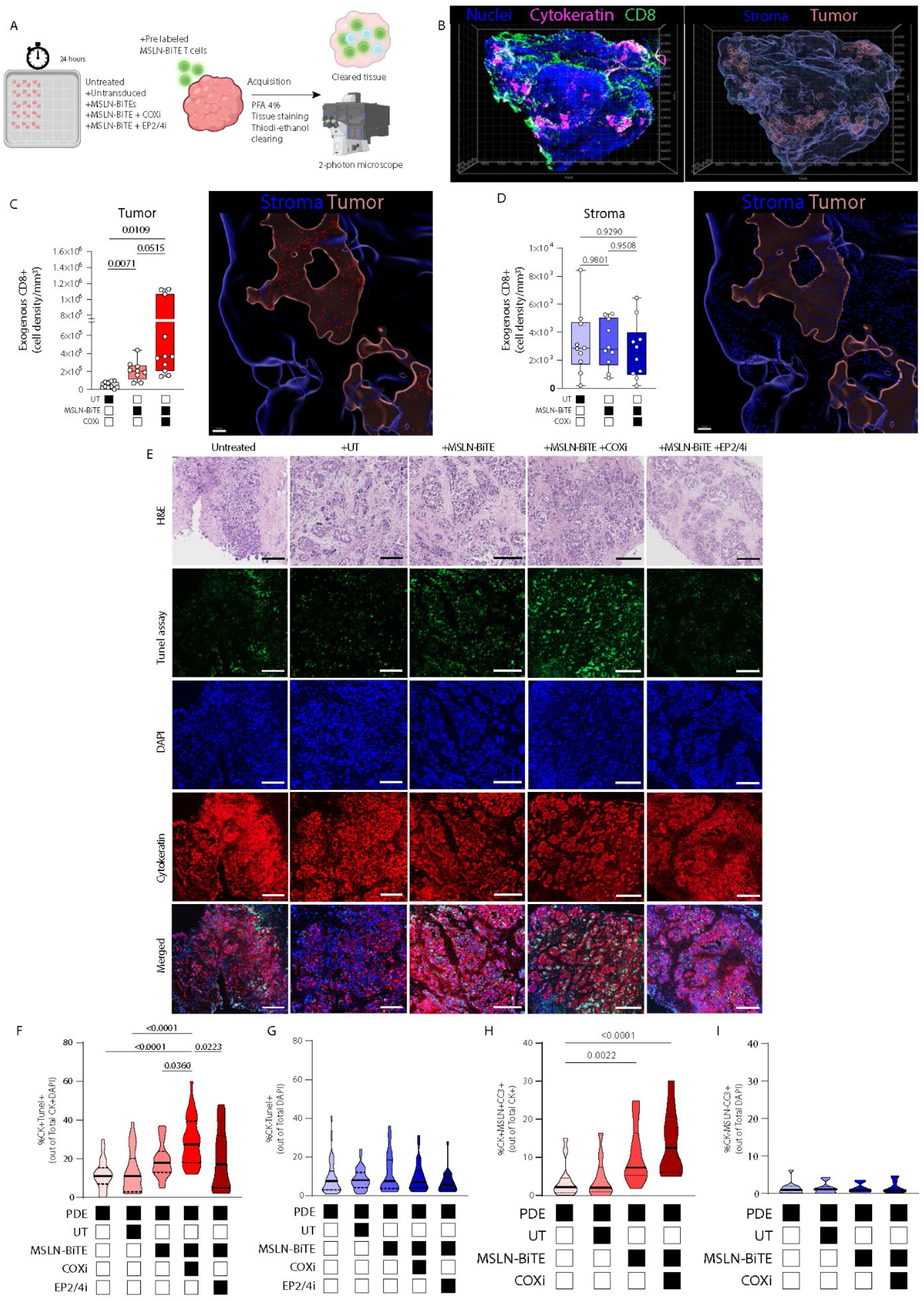
COXi enhances intratumoral accumulation of MSLN-BiTE T cells and increases tumor-islet-restricted DNA damage in PDEs. **A**, Schematic of the volumetric two-photon imaging pipeline used to quantify intratumoral localization of fluorescently labeled *exogenous* T cells in PDEs after 24 h co-culture (UT T cells, MSLN-BiTE, MSLN-BiTE +COXi, or MSLN-BiTE +EP2/EP4i), followed by fixation, pan-cytokeratin staining to define tumor epithelium, DAPI counterstaining, tissue clearing, and two-photon acquisition. **B,** Representative 3D reconstructions showing nuclei (blue), cytokeratin-defined tumor epithelium (magenta), and infiltrating CD8⁺ T cells (green), with corresponding tumor and stroma masks generated by cytokeratin-based segmentation. **C,** Boxplots quantifying *exogenous* CD8⁺ T cell density within the CK⁺ tumor compartment and CK⁻ stromal compartment. **D,** Normalized to imaged surface area, with representative segmentation overlays. **E,** Representative H&E and IF images of DNA damage after 24 h co-culture, showing TUNEL (green), DAPI (blue), cytokeratin (red), and merged channels across conditions. **F-G**, Violin plots showing the fraction of TUNEL⁺ cells among CK⁺DAPI⁺ epithelial cells and among CK⁻DAPI⁺ non-epithelial cells. **H-I,** Violin plots quantifying cleaved-caspase-3 (CC3) positivity within CK⁺MSLN⁺ epithelial cells and within CK⁻ cells. Each dot represents an individual PDE fragment. Statistical analyses were performed using one-way ANOVA (GraphPad Prism v10.1.2); exact P values are indicated on the plots. Figure generated with BioRender. Error bars indicated ± SEM.

We then performed 3D digital reconstruction of the imaged PDEs and segmented tumor and stromal compartments based on CK positivity to precisely localize infiltrating T cells within the PDEs (**Fig. 5B**). Volumetric depth profiling revealed that MSLN-BiTE T cell treatment promoted active CD8⁺ T cell migration across the tumor-stroma interface and deep penetration into CK⁺ tumor nests, with CD8⁺ cell density progressively increasing along the z-axis toward tumor cores, a pattern further accentuated by COX/PGE_2_ inhibition (**Fig. 5C** and **Fig. S5A**).

Quantitative volumetric analysis demonstrated that PDEs treated with MSLN-BiTE T cells displayed dense CD8⁺ T cell infiltration (Celltrace Far Red labeled (T cells, green) cells /total surface area)) and localization within the tumoral compartment (CK^+^DAPI^+^) (**Fig. 5B-C**) with minimal localization in the stromal regions (**Fig. 5D**). UT T cells had similar CD8^+^ T cell densities in tumor and stromal regions concordant with our previous results (**Fig. S4A).** Interestingly, we observed that COX/PGE_2_ axis inhibition further increased the localization of CD8 T cell in CK⁺ tumor areas (**Fig. 5C, bar plot**). Our volumetric data suggests that PGE_2_ may impair T cell motility and demonstrate that COX/PGE_2_ inhibition increases intratumoral CD8⁺ motility and engagement of MSLN-BiTE with tumor cells within the PDEs.

We next asked whether CD8^+^ infiltration and localization in tumor compartments translated into T cell engagement by measuring DNA damage using TUNEL assay by IF. After coculture of PDEs with MSLN-BiTE T cells (± COX inhibition or EP2/EP4 antagonism), tissues were fixed, paraffin-embedded, and sectioned. We then performed TUNEL to detect DNA fragmentation by immunofluorescence, with pan-CK to delineate tumor epithelium and DAPI as nuclear counterstain. Quantitatively, TUNEL staining (green) increased prominently within cytokeratin nests (panCK, red) when PDEs were cocultured with MSLN-BiTE T cells (**Fig. 5E**). The percentage of CK⁺ cells that were TUNEL⁺ further increased with combinatorial MSLN-BiTE and COXi treatment (**Fig. 5F**). By contrast, EP2/EP4 co-treatment produced modest gains, similar to MSLN-BiTE T cells alone, consistent with the superior efficacy of upstream COX inhibition (**Fig. 4F-H**). Of note, the TUNEL⁺ fraction of CK⁻DAPI⁺ cells were not affected by MSLN-BiTE T cells (**Fig. 5G**), indicating that DNA damage was concentrated within tumor epithelium rather than broadly induced in non-tumoral compartments. These data suggest that MSLN-BiTE T cells only exert their tumor reactivity on target and can spare the stromal compartment or healthy cells from cytotoxicity, validating the high specificity of the targeted cells.

Because TUNEL labels DNA strand breaks that can also arise from non-apoptotic DNA damage, we also validated tumor cell apoptosis using a cleaved-caspase 3 (CC3) immunostaining as a more apoptosis-proximal readout, which staining results showed a similar pattern as the TUNEL assay (**Fig. 5H** and **Fig. S5B)**. Specifically, we saw that in contrast to UT T cells, MSLN-BiTE T cells increased CC3 expression in MSLN^+^CK^+^DAPI^+^ (green) cells within the PDEs 24 hrs post coculture which was further augmented by inhibition of COX enzymes. Again, normal cells of the stroma were spared from apoptosis **(Fig. S5B** and **fig. 5I)**.

Together, volumetric two-photon imaging and apoptosis mapping demonstrate that MSLN-BiTE T cells actively penetrate tumor islets, localize within epithelial nests, and induce target-restricted cell death without bystander damage. Upstream COX/PGE_2_ inhibition further intensifies this spatially confined recruitment-and-kill program by enhancing CD8⁺ motility, deepening intratumoral penetration, and coordinating pro-inflammatory myeloid and chemokine networks within intact human TMEs.

### An *ex vivo* response score identifies TME biomarkers for MSLN-BiTE T cell efficacy across NSCLC and HGSOC

Having established that MSLN-BiTE T cells achieve *bona fide* intratumoral localization and tumor-restricted killing, we next sought to integrate cellular activation and soluble immune programs into a unified quantitative framework to compare *ex vivo* responses across NSCLC and HGSOC cohorts.

We therefore derived an *ex vivo* response score anchored on tumor-reactive T cell activation^32^. Antigen-specific engagement was captured by induction of CD8⁺CD137⁺ T cells following exposure to secreted MSLN-BiTEs, quantified for each PDE. Lesions exhibiting a CD8⁺CD137⁺ fold change <2 relative to UT T cell controls were classified as Non-Responder (NR), irrespective of cytokine modulation (**Methods**).

Application of this response score to the NSCLC PDE cohort stratified patients into Responder (R) and NR groups (**Fig. 6A-E**). Using a significant CD137⁺ induction in CD8⁺ T cells as the mandatory criterion, responsive PDEs met the activation threshold and concomitantly exhibited robust increases in cytotoxic mediators including granzyme B, IFNγ, TNFα. In contrast, NR PDEs, defined by less than 2-fold CD137⁺ induction, failed to mount consistent or coordinated immune responses across these readouts (**Fig. 6B-E**).

**Figure 6:**
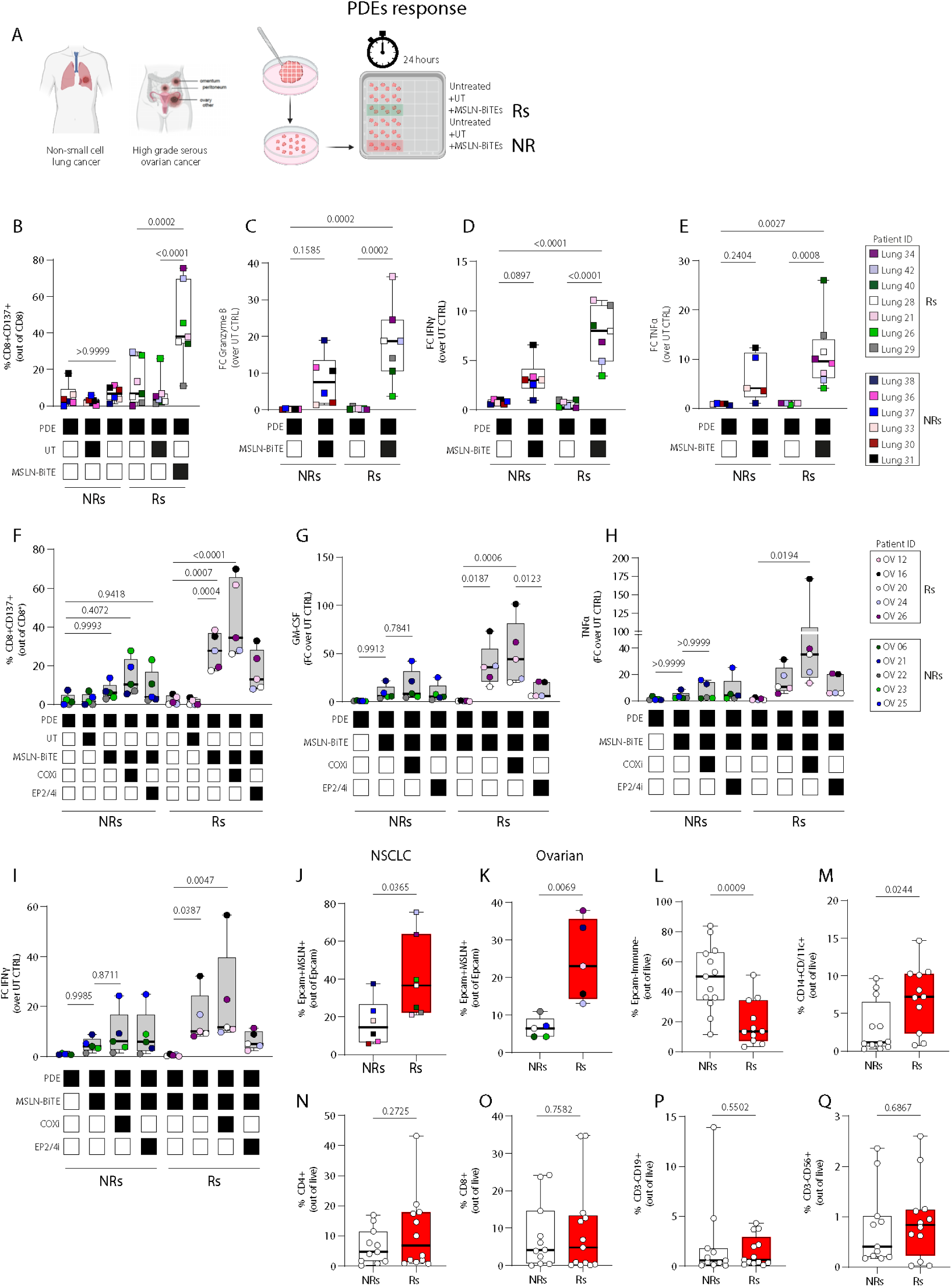
*Ex vivo* response stratification distinguishes MSLN-BiTE responder and nonresponder lesions and reveals baseline features associated with nonresponse. A, Schematic of the PDE coculture assay and response stratification workflow used to classify lesions as responders (Rs) or nonresponders (NRs) after 24-hour exposure to MSLN-BiTE-secreting T cells relative to (UT) T cell CTRL, based on CD8^+^CD137^+^ induction. (**B-E**) NSCLC cohort. (**B**) Induction of CD8^+^CD137^+^ cells among CD8^+^ T cells. (**C-E),** Fold change (FC) over UT control for granzyme B (GZMB; C), IFNγ (D), and TNFα (E) in Rs versus NRs after MSLN-BiTE coculture. **F-I,** HGSOC cohort. **F**, Induction of CD8^+^CD137^+^ cells among CD8^+^ T cells across the indicated treatment conditions (UT, MSLN-BiTE, MSLN-BiTE +COXi, and MSLN-BiTE +EP2/EP4i). **(G-I)** Corresponding modulation of GM-CSF (**G**), TNFα (**H**), and IFNγ (**I**), shown as FC over UT control, in Rs versus NRs. (**J-K)** Baseline target availability, quantified as the percentage of EPCAM^+^MSLN^+^ tumor cells among EpCAM^+^ cells, in NSCLC (**J**) and HGSOC (**K**) Rs versus NRs. **L-Q,** Baseline tissue composition according to response class, including stromal cells (EPCAM^-^immune^-^ live cells; **L**), CD14^+^CD11c^+^ myeloid cells (**M**), CD4^+^ T cells (**N**), CD8^+^ T cells (**O**), CD19^+^ B cells (**P**), and CD3^-^CD56^+^ NK cells (**Q**). In **B-I**, each point represents an individual explant fragment and is colored by patient; condition matrices are shown below each panel. In **J-Q**, each point represents one patient. Box plots show the median and interquartile range. Exact *P* values are indicated in the panels. Figure created with BioRender.

Overall, 7 of the 13 NSCLC patients’ PDEs qualified as R, while the remainder clustered as NR, underscoring marked inter-patient heterogeneity in functional sensitivity to MSLN-BiTE T cell therapy.

We next applied the same response score to the spatially profiled HGSOC PDE cohort. Among the 10 OC patients analyzed, 5 were classified as R and 5 as NR, revealing a near-even distribution of functional sensitivity to MSLN-BiTE therapy that closely mirrored the NSCLC cohort (**Fig. 6B-E**). Consistent with the scoring criteria established in NSCLC, R HGSOC lesions displayed significantly higher frequencies of activated CD8⁺CD137⁺ T cells upon MSLN-BiTE exposure compared with NR (**Fig. 6F**).

Functional stratification was further supported by robust induction of inflammatory mediators, including GM-CSF, TNFα, and IFNγ, in Rs lesions treated with MSLN-BiTEs (**Fig. 6G-I**). Notably, while combined MSLN-BiTE and COX inhibition did not further increase CD8⁺CD137⁺ T cell frequencies, COXi selectively amplified GM-CSF and TNFα production, indicating coordinated engagement of myeloid and antigen-presenting cell-associated inflammatory programs downstream of T cell activation in R (**Fig. 6G-H**). These findings suggest that COX inhibition primarily augments the quality and tissue-level amplification of immune responses.

Importantly, combinatorial MSLN-BiTE and COX inhibition rescued functional responses in two out of 5 NRs PDEs (**Fig. 6F**), indicating that prostanoid signaling represents a dominant albeit not exclusive axis of TME-mediated resistance. The persistence of non-responsiveness in a subset of lesions despite COX blockade suggests the coexistence of additional mechanisms that constrain immune activation.

Importantly, R and NR stratification could be explained by differences in baseline antigen availability, stromal dominance, and pre-existing myeloid context. Specifically, the fraction of MSLN⁺ tumor cells (out of EPCAM⁺) was significantly higher in responsive lesions in both NSCLC and HGSOC cohorts (**Fig. 6J-K**), indicating that reduced target density contributes to limited BiTE engagement in NR. In parallel, non-responsive lesions exhibited a significantly higher proportion of Epcam⁻immune⁻ stromal cells, consistent with a stromally dominant tissue architecture (**Fig. 6L**), whereas R were enriched for CD14⁺CD11c⁺ myeloid cells, suggestive of greater antigen-presenting and immune-licensing capacity (**Fig. 6M**). By contrast, baseline frequencies of bulk lymphoid populations, including CD8⁺ and CD4⁺ T cells, B cells, and NK cells (CD3⁻CD56⁺), were comparable between groups (**Fig. 6N-Q**).

Together, these data indicate that sparser antigen availability combined with stromal enrichment and limited myeloid licensing constitutes a major resistance axis to MSLN-BiTE efficacy, whereas lymphocyte abundance per se is not a determining factor.

### Spatial transcriptomics reveals response-specific immune hubs and stromal barrier states in non-responder lesions

To investigate whether *ex vivo* functional responses correspond to localized transcriptional immune remodeling within intact TME, we performed spatial transcriptomic profiling of OC PDEs treated with MSLN-BiTE T cell alone or in combination with COXi inhibition. This analysis was designed to determine whether the baseline features associated with resistance, namely stromal dominance and limited myeloid licensing capacity, manifest as spatially organized transcriptional states within intact tissue, and whether responder lesions assemble distinct immune-permissive niches upon treatment.

Spatial transcriptomic data were generated from five PDEs derived from four HGSOC patients (post treatment with MSLN-BiTE T cells alone or in combination with COX inhibition), including both R and NR lesions as defined by our *ex vivo* response score. Spots were deconvolved to refined immune and stromal states using an already published dataset (**Fig. S6A**), and then segmented into tumor islets, stroma, and mixed rims, enabling spatially resolved analysis of treatment-induced transcriptional changes (**Fig. 7A-C**). This approach ensured that cell-state, pathway and gene readouts were interpreted within their precise anatomical context.

**Figure 7:**
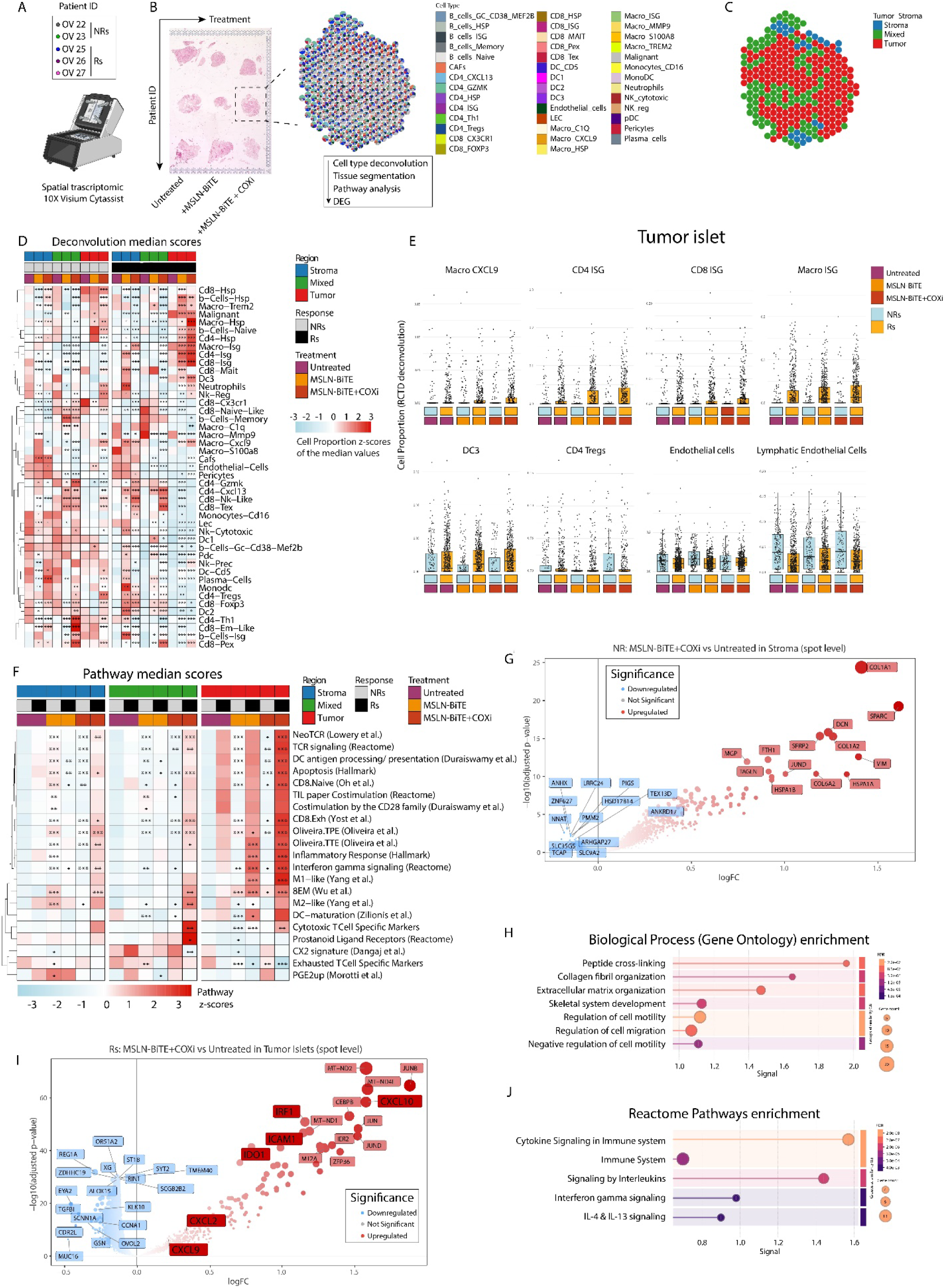
Spatial transcriptomics identifies responder-specific immune hubs in HGSOC PDEs treated with MSLN-BiTE with or without COX inhibition. **A,** Overview of the spatial transcriptomics workflow (10x Genomics Visium CytAssist) and the HGSOC PDE set analyzed, including R and NR lesions as defined by the *ex vivo* response score. **B,** Representative H&E images and corresponding Visium capture areas across treatment conditions (Untreated, +MSLN-BiTE, +MSLN-BiTE + COXi), with the downstream analysis steps indicated (tissue segmentation, cell-type deconvolution, pathway scoring, and differential expression). **C,** Spatial annotation of Visium spots into tumor, stroma, and mixed regions based on tissue segmentation. **D,** Heatmap of median cell-state deconvolution scores (z-scored) across samples, stratified by anatomical region, response class (Rs vs NRs), and treatment condition. **E,** Spot-level deconvolution proportions for selected immune and stromal states within tumor islets, grouped by treatment and response class. **F,** Heatmap of median pathway scores (z-scored) across regions, response class, and treatment. (**G-H**) NR stroma differential expression for MSLN-BiTE + COXi versus Untreated at spot level: (**G**) volcano plot with representative genes highlighted in NRs stroma compartment. (**H**) Gene Ontology biological process enrichment of upregulated genes. (**I-J**) Rs tumor-islet differential expression for MSLN-BiTE + COXi versus Untreated at spot level. (**I**) Volcano plot with representative genes highlighted in responders’ tumor islet; (**J**) Reactome pathway enrichment of upregulated genes. Figure generated with BioRender.

Cell dtype deconvolution revealed a clear divergence between R and NR lesions following treatment (**Fig. 7D**). In R lesions, particularly under MSLN-BiTE plus COX inhibition, tumor islets and adjacent mixed regions were enriched for interferon-stimulated T cell states (*CD8-ISG*, *CD4-ISG*) and inflammatory, interferon-licensed macrophages (*Macro-CXCL9*, *Macro-ISG*), alongside dendritic cell populations (*DC3*) associated with antigen processing and presentation. These immune-enriched states were accompanied by a contraction of stress-dominated programs, including heat-shock-associated T cell and macrophage states (*CD8-HSP*, *Macro-HSP*). Notably, these changes were spatially confined to tumor-containing compartments rather than diffusely distributed across the stroma (**Fig. 7D**).

In contrast, NR lesions exhibited a distinct spatial organization characterized by the persistence of stromal-associated non-immune populations, including lymphatic endothelial cells (LECs), vascular endothelial cells, and cancer-associated fibroblasts (CAFs), which remained dominant across treatment conditions (**Fig. 7D, E**). These populations were preferentially localized to stromal regions and mixed rims and showed limited modulation following MSLN-BiTE or MSLN-BiTE plus COXi treatment. Consistent with the FACS-based stratification (**Fig. 6L)**, spatial transcriptomic quantification revealed that NR PDEs were characterized by a higher stromal tissue fraction, whereas R PDEs tended to display reduced stromal representation and a relative expansion of tumor islet areas, although differences in mixed and tumor compartments did not reach statistical significance (**Fig. S6C**).

Pathway-level analyses reinforced these spatial distinctions (**Fig. 7F**). In R lesions, treatment induced stepwise increases, most pronounced with MSLN-BiTE plus COXi, in IFNγ signaling, inflammatory response, T cell receptor signaling and costimulation, antigen processing and presentation, and apoptosis (**Fig. 7F**). These shifts were concentrated within tumor islets and mixed regions, consistent with the formation of localized immune activation hubs. Indeed Supplementary **Fig. S6D**, **E**, and **F** demonstrate that R lesions exhibit treatment-induced IFNγ pathway activation that intensifies across successive peritumoral layers toward tumor islets and adjacent mixed regions, while NR lesions show blunted or absent IFNγ responses across all spatial layer (**Fig. S6D-F**).

Transcriptionally, NR stroma was enriched for programs linked to extracellular matrix organization, collagen fibril assembly, regulation of cell motility, vascular organization, and tissue remodeling, consistent with a structurally reinforced microenvironment (**Fig. 7G-H**). Importantly, these stromal programs were not accompanied by induction of immune chemokines, antigen-presentation pathways, or interferon-responsive modules (**Fig. S6G-H**), consistent with the absence of coordinated cytokine and activation signatures observed by FACS in NR PDEs (**Fig. 6E-H**), suggesting a barrier-like phenotype that may constrain immune cell recruitment and effector engagement.

Furthermore, in NR lesions, immune and apoptotic pathways showed minimal induction, while matrix remodeling, stress response, and structural maintenance programs remained prominent. Where prostanoid-related signatures were detectable, they trended downward with COX inhibition but did not translate into coordinated immune activation in NR lesions (**Fig. 7F**).

Tumor-islet restricted differential expression analysis further distinguished the two response states (**Fig. 7I-J, Fig. S6I**). In R lesions, MSLN-BiTE plus COXi treatment induced chemokines and immune interaction genes such as *CXCL9*, *CXCL10*, *ICAM1*, and *IRF1*, consistent with leukocyte recruitment, adhesion, and interferon licensing within epithelial nests, alongside induction of *IDO1* (**Fig. 7I-J**). In contrast, NR lesions preferentially upregulated genes associated with cellular stress and adaptation, including heat-shock chaperones (*HSPA1A*, *HSPA1B*, *HSPB1*, *DNAJB1*), oxidative stress mediators (*SOD2*, *FTH1*), and extracellular matrix components and *VIM* without assembling the IFN-chemokine module observed in responsive ones (**Fig. S6G-H,** **Fig. 7G-H**).

Collectively, these spatial data demonstrate that the pharmacodynamic state defined by the *ex vivo* response score is transcriptionally and anatomically localized to tumor islets in responsive lesions providing a spatial correlation with CD137⁺ activation, cytokine induction, and myeloid engagement. In these regions, MSLN-BiTE initiates, and COX inhibition amplifies, an interferon-linked, antigen-presentation-competent and apoptosis-associated immune program. In contrast, NR lesions remain dominated by stromal barrier states characterized by LECs, endothelial cells, and CAF-driven remodeling programs and fail to assemble coordinated immune hubs under identical conditions. These findings underscore the central role of spatial TME organization in shaping responsiveness to BiTE-based immunotherapy and highlight stromal remodeling as a key axis of resistance.

## Discussion

Resistance to ACT in solid tumors is determined within native tissue, where stromal architecture, myeloid circuits, and locally produced suppressive mediators jointly shape T-cell recruitment, activation, and persistence. This creates a modeling gap, because many commonly used systems disrupt spatial neighborhoods and soluble gradients that control effector implementation. Here, we establish a human-based NAM for ACT pharmacodynamics and stratification that bridges reductionist 3D epithelial models and organotypic PDEs to interrogate MSLN-redirected BiTE-secreting T cells within preserved tumor-stroma structure, resident immune infiltrates, and endogenous immunosuppressive programs.

A key insight from the PDE platform is that redirected T cell activity is heterogeneous and quantitatively constrained across lesions. Compared with antigen-saturated epithelial models, PDEs reveal native ceilings imposed by variable target density together with intact microenvironmental suppression. While spheroids and PDTs are informative for benchmarking antigen dependence under 3D geometry, they lack stromal and myeloid compartments required to capture tissue-level licensing and recruitment loops. In PDEs, responses are instead reflected by coupled cellular and soluble readouts, including CXCL9/10 and GM-CSF, which report on interferon-licensed, APC-associated amplification beyond tumor-intrinsic killing.

Within this intact context, MSLN-BiTE T cells consistently elicited antigen-dependent responses across NSCLC and HGSOC PDEs, marked by induction of CD8⁺ activation, inflammatory cytokines, and CXCR3 ligands. BiTEs acted as “diffusible bridges” by engaging both engineered and bystander/endogenous T cells. These responses were spatially resolved: volumetric two-photon imaging demonstrated deep CD8⁺ accumulation within CK⁺MSLN⁺ epithelial nests, and apoptosis mapping revealed tumor-restricted cell death with no impact on non-tumoral compartments. These findings establish that redirected BiTE activity can be executed with high spatial specificity in intact human tissue, consistent with diffusible CD3ε engagement that recruits proximate endogenous T cells and extends cytotoxic reach beyond direct engineered-cell contacts.

These spatial readouts have direct translational relevance. Human tumors span immune-inflamed, immune-excluded, and immune-desert phenotypes that are often inferred from planar CD8⁺ immunohistochemistry, which under-evaluates ITH and spatial distribution. By enabling three-dimensional, compartment-resolved quantification of immune cell localization, PDE volumetrics provide an integrated assessment of whether effectors penetrate CK⁺ tumor nests or remain stromal-restricted. Within this framework, PDEs support two complementary applications: testing TME-targeted interventions in immune-infiltrated or partially excluded lesions, and simulating ACT infusion in immune-poor lesions to directly measure infiltration and tumor-islet killing under preserved stromal constraints

Spatial transcriptomics further refined this interpretation by localizing response programs to discrete tissue domains. In R lesions, tumor islets and adjacent mixed regions assembled immune hubs marked by interferon signaling, antigen processing and presentation, costimulatory pathways, dendritic cell maturation, and cytotoxic programs. The emergence of responder-specific immune hubs enriched for interferon-licensed macrophage and dendritic cell states is also consistent with the view that pre-existing myeloid-T cell network architecture conditions ACT efficacy^26, 28, 33^. In contrast, NR lesions were dominated by tumor associated vasculature, ECM remodeling and EMT-associated-programs, with limited induction of interferon-chemokine modules. This dichotomy indicates that ACT resistance can be encoded in spatial tissue states that preclude productive immune implementation.

A central mechanistic finding is that the COX/prostanoid axis quantitatively constrains ACT efficacy in intact human tissue. Baseline PGE_2_ burden inversely correlated with CD8⁺CD137⁺ activation and granzyme B release after MSLN-BiTE T cell exposure, defining a prostanoid-dependent ceiling on cytotoxic amplification consistent with our prior work^16, 17^. Pharmacologic COX inhibition reduced PGE_2_, amplified activation of engineered and endogenous CD8⁺ T cells, and shifted the myeloid compartment toward a CD68⁺CD80⁺ inflammatory, APC-supportive phenotype. COX blockade intensified an IFNγ/CXCL9/10 axis integrating T cell and myeloid programs, translating into deeper CD8⁺ penetration of CK⁺MSLN⁺ nests and increased epithelial apoptosis without detectable bystander toxicity. Selective EP2/EP4 antagonism produced weaker effects, consistent with partial relief of prostanoid signaling compared with upstream COX inhibition.

Beyond mechanism, the PDE platform enables functional stratification and identification of resistance archetypes. Integrating cellular activation, soluble mediators, spatial imaging, and transcriptional programs segregated R and NR lesions according to both effector competence and TME-imposed constraints. Prostanoid burden emerged as a candidate biomarker for ACT responsiveness and selection of COX axis combinations, whereas NR lesions enriched for ECM/EMT programs linked to immune exclusion. These findings suggest that PDEs can inform rational combination strategies targeting orthogonal stromal resistance circuits not addressed by antigen redirection alone. Although MSLN-redirected BiTE-secreting T cells were used here as a clinically relevant ACT model in NSCLC and HGSOC, the PDE platform is inherently antigen-agnostic and compatible with diverse immune checkpoint inhibitors and ACT modalities including TIL products, CAR-T cells, TCR-engineered T cells, and can be readily redeployed against various antigen targets (e.g., FOLRA^34^, CLDN6^35^, CLDN18.2^36^) as they mature^37^. Our results support a model in which three-dimensional tissue architecture imposes shared constraints on immune cell infiltration and activation across ACT platforms, highlighting tissue-encoded resistance rather than effector design or antigen selection, as a primary determinant of functional efficacy in solid tumors.

This study has limitations inherent to *ex vivo* systems. PDEs capture tumor-local pharmacodynamics and tissue-restricted killing, not systemic exposure, vascular delivery, or whole-body cytokine dynamics. Lack of perfusion precludes pharmacokinetic inference, and dosing reflects local target engagement. Nevertheless, PDEs preserve resident myeloid architecture and licensing states that can condition early ACT implementation *in situ*^26, 28, 33^.

In summary, this work establishes PDEs as a physiologically grounded platform for interrogating adoptive cell therapy within conserved human TMEs and identifies the COX/prostanoid axis as a tunable, tissue-imposed constraint on MSLN-BiTE-driven immunity. By linking antigen-dependent activation to spatially organized immune hubs and revealing a modifiable prostanoid brake, our study provides a framework for mechanism-based patient stratification and rational combination design in solid-tumor ACT. More broadly, these findings underscore that the success of adoptive immunotherapy is governed not only by effector cell design, but by the spatial and biochemical constraints encoded within intact human tissues, arguing for a shift toward tissue-preserving preclinical models that capture early pharmacodynamic responses under native conditions. Consistent with recent spatial dissection of prostaglandin-mediated immune suppression^14^, our data support a model in which ACT efficacy is governed by the ability to overcome regionally encoded biochemical barriers that restrict interferon signaling and immune hub formation within tumor-proximal tissue.

## Acknowledgments

We are grateful to the patients and their families for their dedicated collaboration.

We thank Jean-Paul Rivals, Marzio Bergomi, Audrey Goupil and all the team from CHUV Biobank, Center of Experimental Therapeutics (CTE) for their assistance.

We thank the Medical Oncology Service, Department of Oncology, the Institute of Pathology and the Pathology biobank, the Unit of Translational Onco-pathology and the Department of Obstetrics and Gynecology, Lausanne University Hospital (CHUV), Switzerland.

We thank the Division of Thoracic Surgery, Department of Surgery, Lausanne University Hospital (CHUV), Switzerland.

We thank Véronique Noguet-Broguier, Luigi Bozzo and Alexandre Benechet from the histopathology and Cellular Imaging facilities for technical support, respectively of the Faculty of Biology and Medecine (FBM), University of Lausanne, Switzerland.

We thank the Lausanne Genomic Technologies Facility for RNA-seq analysis.

We thank the Bioengineering and Organoids Technology Platform (BET) of École Polytechnique Fédérale de Lausanne (EPFL) for assistance with the Cell Discoverer 7 imaging system.

## Funding resource

This work was supported by the ISREC foundation, the Ludwig Institute for Cancer Research (Myeloid Cells in Cancer Initiative, MCCI project; LAbCore), the DOD OCA Early Career Investigator (ECI) W81XWH2210703 Award OC210038 to D.D.L.

## Materials availability

The anti-MSLN 13F08 BiTE is under development for ACT applications. Its sequence is proprietary to the UNIL-CHUV LAbCore Platform and is not disclosed in this study. Requests for further information should be addressed to S.D.

## Materials and methods

### Ethics approval

This study received central approval by Lausanne University (UNIL) and Ludwig Cancer Research Lausanne Branch institutional review board. All procedures were performed according to the Declaration of Helsinki guidelines. These specimens were collected under approved translational protocol and informed consent (2016-02094, Pre-IT/ CHUV_DO_pre-IT_2016) and managed by the Centre de Thérapies Experimentales (CTE) of CHUV’s Department of Oncology. Specimens were coded (de-identified) and stored at CTE’s biobank (BBO-CTE - AO_2021_00011), then transferred to the Dangaj laboratory for analyses using three translational research (2017-00359, 2017-00305 and 2017-00490).

### Tumor sample processing

Solid tumor lesions were macroscopically selected by a pathologist from the resected tumor material and part of the tumor was collected in R10 (RPMI 1640 medium (Thermo Fisher Scientific, 61870-010) supplemented with 10 % FBS (Thermo Fisher, A5256701), 100 U/mL penicillin, and 100 µg/ml streptomycin sulfate (BioConcept, 4-01F00-H) for subsequent PDE cultures. A second part of the tumor was embedded in paraffin for histological analysis and for analysis of tumor cell content and architecture within the collected lesion. Tissue materials collected for subsequent PDE cultures were immediately processed by manual cutting into small tumor fragments PDE of 2 mm^3^ size on ice. After processing, a number of single PDEs from different regions within a tumor were mixed to ensure uniform representation of the tumor lesion and were frozen in cryovials containing 1 ml of FBS with 10% dimethylsulfoxide (DMSO) (PanReac AppliChem, A3672.0100) with 8-12 PDEs per vial.

### Cell line

Human OVCAR-8 (RRID:CVCL_1629) cell line was a generous gift from Steven Dunn’s lab coming from UPENN Ovarian Cancer Research Center and used as a model tumor cells for studying meso-CAR-T and BiTE therapy. Cells were maintained in R10. All cells were maintained at 37°C, 5% CO_2_ in a humidified incubator and medium was renewed 3 times per week to ensure exponential growth conditions.

### MSLN-CAR and MSLN-BiTE T cell products

The fully-human anti-MSLN 13F08 CAR and T cell-secretory BiTE warhead was developed using an in-house in vitro discovery platform (SD). Primary human CD4^+^ and CD8^+^ T cells were isolated from peripheral blood mononuclear cells (PBMCs) obtained from anonymous healthy donors (buffy coats, Transfusion Interrégionale CRS) using Ficoll-Paque™ density gradient centrifugation (Ficoll-Paque™ PLUS, Cytiva) followed by immunomagnetic negative selection (EasySep™ Human CD4^+^ and CD8^+^ T Cell Isolation Kits, Stem Cell Technologies) according to the manufacturer’s instructions.

### Virus production

Retroviral supernatant was generated by triple transfection of HEK293T cells (ICLC Cat# HTL04001, RRID:CVCL_0063). HEK293T cells at 80% confluence in T150 flasks were co-transfected with 18 μg Gag-Pol (Peq-Pam), 7 μg RD114, and 22 μg of the retroviral vector encoding the gene of interest using TurboFect Transfection Reagent (ThermoFisher Scientific). Plasmids were mixed with 3 mL Opti-MEM™ I Reduced Serum Medium (Gibco) and 120 μL TurboFect, then incubated for 30 min at room temperature before addition to the cells. After incubation, cultures were returned to the incubator, and the medium was replenished the following day. At 48 hrs and 72 hrs post-transfection, viral supernatant was collected, filtered through a 0.45 µm filter (Sartorius), and concentrated by ultracentrifugation at 24,000 g at 4°C for 2 hrs (Avanti JXN-30 high-speed centrifuge). Virus was used immediately or aliquoted, frozen on dry ice, and stored at -80°C. Lentivirus was produced and concentrated using the same protocol, substituting the plasmid mixture with 18 μg pCMVR8.74 (RRID:Addgene_22036), 7 μg VSV-G, and 15 μg lentiviral vector encoding the gene of interest.

### T cell transduction

Freshly isolated human CD4^+^ and CD8^+^ T cells were activated on day 0 using CD3/CD28 human T-Activator Dynabeads™ (ThermoFisher Scientific) at a 2:1 bead-to-cell ratio and cultured in R10 supplemented with 50 IU/mL recombinant human IL-2 (PeproTech) at 1×10^6^ cells/mL (1 mL per well) in 24-well plates. T cell transductions were performed on day 1 post-activation. Retroviral transductions utilized RetroNectin (Takara Bio)-coated, non-treated tissue culture plates (Greiner Bio-One) following the Takara protocol. Virus binding to RetroNectin was achieved by centrifuging the plates for 1.5 hrs at 32°C at 2000 g according to the manufacturer’s instructions. After centrifugation, activated donor-matched CD4^+^ and CD8^+^ T cells were pooled at 20% CD4^+^ and 80% CD8^+^ and added to the RetroNectin/retrovirus-coated plates. For 24-well plates, 1×10^6^ T cells were added per well; for 48-well plates, 0.5×10^6^ T cells were added. Plates were centrifuged at 300 g for 10 min to promote T cell contact with viral particles, then returned to the incubator. Lentiviral transductions were performed by directly adding viral particles to activated T cells (pooled donor-matched CD4^+^ and CD8^+^ at 20% and 80%, respectively). From day 2 post-transduction, T cells were expanded in complete RPMI medium supplemented with human IL-7 and IL-15 cytokines (each at 10 ng/mL; Miltenyi Biotec) and maintained at 0.5×10^6^ cells/mL as previously described to ensure optimal T cell growth. Dynabeads™ were removed on days 5-6, and transduction efficiency and T cell viability were assessed by flow cytometry on day 7 unless otherwise specified.

### Extracellular and Intracellular staining of transduced T cells

Transduction efficiency of Mesothelin CAR and Mesothelin BiTE T cells was evaluated by cell-surface staining of epitope tags using anti-cMyc antibody (Creative Diagnostics Cat# DMAB8571, RRID:AB_2392346). The anti-DYKDDDDK (FLAG) Epitope Tag Alexa Fluor 647 antibody (R&D Systems Cat# IC8529R, RRID:AB_3656740) was used to detect FLAG-tagged 13F08 BiTE (MSLN-BiTE). After cell surface staining, cells were subjected to intracellular staining as previously described to detect expressed BiTEs. Cell viability was determined using LIVE/DEAD UV Zombie. Flow cytometry analysis was performed using a CytoFLEX instrument (Beckman Coulter).

Both MSLN-CAR and BiTE T cells were cryopreserved in FBS 10 % DMSO for further use.

### OVCAR-8 spheroid culture

Spheroids were generated using matrix-free 96 well microwell plates (SunBioscience) designed to promote homogeneous three-dimensional aggregation. Single-cell suspensions were seeded at 10,000 cells per well in 30 µL of culture medium and allowed to sediment for 30 min at 37 °C under 5% CO_2_ to facilitate uniform spheroid formation within the microcavities. Wells were subsequently overlaid with 170 µL of R10. Spheroids were maintained under standard culture conditions with medium replacement every 2-3 days and were used for downstream experiments once a diameter of 200-300 µm was reached.

### Patient derived tumoroids culture

Tumor tissue was obtained either by core needle biopsy or by surgical resection and processed for tumoroid culture immediately after surgery or from cryopreserved material. Small biopsy specimens were subjected to fine-needle mechanical dissociation using a 25-gauge needle, following previously published protocols, generating small tissue fragments that were directly embedded in Matrigel (Corning, 356231) and overlaid with growth factor-supplemented medium. Tumor tissue derived from surgical resections was minced into small pieces and enzymatically digested using an in-house cocktail containing collagenase I (50 U/mL, Gibco), collagenase IV (50 U/mL, Gibco), DNase I (30 U/mL, Sigma-Aldrich), and RNasin (1:1,000, Promega), followed by mechanical dissociation and filtration through a 70 µm cell strainer to obtain a single-cell suspension. Single cells were embedded in extracellular matrix at a density of 50,000 cells per 30 µL drop in a mixture of 80% Matrigel and 20% growth factor-enriched medium; after polymerization for 30 min at 37 °C, droplets were overlaid with histotype-specific growth factor-supplemented medium (lung or ovarian), which was replaced every 2-3 days.

Tumoroids were passaged approximately every two weeks by mechanical disruption of Matrigel droplets in cold PBS followed by incubation for 1 h at 4 °C in Cell Recovery Solution (Corning) to remove residual matrix, and cells were re-embedded in fresh extracellular matrix. Tumoroids could be cryopreserved after matrix removal in FBS 10% DMSO. Tumoroid growth and morphology, including the presence of necrotic cores, were routinely monitored by brightfield microscopy, and cultures were passaged when tumoroids reached a diameter of 200-300 µm, typically within 10-14 days.

### Tumoroids culture media

For lung tumoroids, cultures were maintained in Advanced DMEM/F-12 medium supplemented with GlutaMAX (Gibco, 10565018), 10 mM HEPES (Gibco, 15630080), and 100 U/mL penicillin-streptomycin (Gibco, 15140122). The medium was further supplemented with 1× N21-MAX Supplement (R&D Systems, AR008), 1.25 mM N-acetylcysteine (Sigma-Aldrich, A9165), 0.5 µM A83-01 (Tocris Bioscience, 2939), 0.5 µM SB202190 (Tocris Bioscience, 1264), 5 mM nicotinamide (Tocris Bioscience, 4106), 5 µM Y-27632 dihydrochloride (Sigma-Aldrich, Y27632), 0.5 µg/mL recombinant human R-spondin 1 (PeproTech, 120-38), 100 ng/mL recombinant human Noggin (PeproTech, 120-10C), 100 ng/mL recombinant human FGF-10 (R&D Systems, 345-FG), and 25 ng/mL recombinant human FGF-7 (R&D Systems, 251-KG).

For ovarian tumoroids, cultures were maintained in Advanced DMEM/F-12 medium supplemented with GlutaMAX (Gibco, 10565018), 10 mM HEPES (Gibco, 15630080), and 100 U/mL penicillin–streptomycin (Gibco, 15140122). This base medium was supplemented with 0.5 µM A83-01 (Tocris Bioscience, 2939), 1× N-2 Supplement (Gibco, 17502-048), 1 mM nicotinamide (Tocris Bioscience, 4106), 10 µM Y-27632 dihydrochloride (Sigma-Aldrich, Y27632), 10 ng/mL epidermal growth factor (Gibco, E9644), and 10 ng/mL recombinant human BMP-2 (Thermo Fisher Scientific, PHC7141).

### *In vitro* coculture

Two days prior to coculture, MSLN-CAR or MSLN-BiTE T cells were thawed and cultured in R10, supplemented with IL-7 and IL-15 at 20 ng/mL each, at a density of 1.5 × 10⁶ cells/mL in 24 well plates to allow functional recovery. On day 2, cytokine withdrawal was initiated by sequential dilution of the culture medium 1:2 with fresh R10, performed twice a day, resulting in T cell conditioning in cytokine free medium prior to coculture. Prior to coculture with patient derived tumoroids, Matrigel was removed as previously described.

Cell lines, patient derived spheroids, and PDTs were cocultured with effector T cells at a 1:1 effector to target ratio in a matrix free setting, either in microcavity plates or in standard 96 well U bottom plates in 200 ul R10 at 37 °C overnight. For confocal microscopy-based assessment of T cell infiltration in spheroids, microcavity plates were used to ensure homogeneous spheroid formation and reproducible replicate size.

Brightfield images were acquired at defined time points using a 5× objective.

### Histological analysis and tissue selection

Patient derived explants were fixed overnight at 4°C in 4% formalin solution (Sigma Aldrich) for standard histopathological processing, followed by paraffin embedding according to routine protocols. Formalin fixed paraffin embedded blocks were sectioned at a thickness of 4 µm using a microtome. Hematoxylin and eosin staining was performed using standard procedures to assess tissue architecture, cellular morphology, and the relative distribution of tumor, stromal, and necrotic areas. Stained sections were used for qualitative histopathological evaluation and for downstream quantitative image analysis.

For quantitative tissue qualification, whole slide images of hematoxylin and eosin stained sections were analyzed in QuPath (v0.5.0). A supervised pixel classification approach was implemented to segment each explant into three compartments, tumor, stroma, and necrosis. Briefly, a subset of representative regions was manually annotated for each class to train the classifier that was then applied to the full tissue section to generate compartment specific masks for tumor, stromal, and necrotic areas. The absolute area of each compartment was quantified within QuPath and normalized to the total tissue area to compute the percentage content of tumor, stroma, and necrosis for each explant.

Explants were selected for downstream functional assays based on predefined quantitative thresholds. Only samples showing a greater than 3> fold enrichment of tumor area relative to both stromal and necrotic compartments were retained, calculated as tumor area divided by stromal area and tumor area divided by necrotic area.

### *Ex vivo* treatment of patient derived tumor fragments

Cryopreserved explants were rapidly thawed and subsequently cut into uniformly sized fragments of approximately 1 to 2 mm³. For each condition, a high number of small fragments, typically 20 to 40 per sample, were randomly distributed in a 96 well membrane bottom plates (Sarstedt) with 200 μL R10 per well. and maintained under gentle rotation.

Where indicated, PDEs were pretreated for 1 to 2 h prior to coculture with either a COXi, ketorolac, at a final concentration of 10 µM, or with an EP2 and EP4 receptor inhibitor at the same concentration. Following pretreatment, explants were directly subjected to coculture with MSLN-BiTE T cells at a 1:1 effector to target ratio under the conditions described above.

### Flow cytometry analysis

Spheroids, PDTs, and PDEs were analyzed by flow cytometry to quantify T cell activation following culture. Tumor tissues were dissociated into single-cell suspensions using enzymatic digestion, as described above, and filtered through a 70 µm cell strainer. Cells were stained with Zombie UV fixable viability dye (BioLegend, 423107) and subsequently incubated with an Fc receptor blocking reagent (Miltenyi Biotec Cat# 130-059-901, RRID:AB_2892112). Surface marker staining was performed using premixed antibody cocktails in PBS 2% FBS and incubated for 15 min at room temperature in the dark prior to downstream acquisition.

T cell activation was assessed using a surface marker panel including anti CD45 (BioLegend, Cat# 304034, RRID:AB_2563465), anti CD3 (BioLegend Cat# 300436, RRID:AB_2562124), anti CD8a (BioLegend Cat# 301042, RRID:AB_2563505), anti CD4 (BioLegend Cat# 317438, RRID:AB_11218995), and anti CD137 (BioLegend Cat# 309808, RRID:AB_394067). Transduced T cells were identified using an anti-FLAG (DYKDDDDK) antibody conjugated to Alexa Fluor 647 (R&D Systems Cat# IC8529R, RRID:AB_3656740).

For PDEs, immune cell composition was additionally characterized using a multiparametric surface marker panel that included anti CD19 (BioLegend Cat# 302216, RRID:AB_314246), anti CD14 (BioLegend Cat# 301820, RRID:AB_493695), anti CD68 (BioLegend Cat# 333808, RRID:AB_1089056), anti CD80 (BioLegend Cat# 305222, RRID:AB_2564407), and anti CD206 (BioLegend Cat# 321136, RRID:AB_2687200). When indicated, tumor cells were identified using anti EpCAM (BioLegend Cat# 324232, RRID:AB_2564301) and anti-MSLN (Santa Cruz Biotechnology Cat# sc-33672, RRID:AB_627930). Unless otherwise specified, all antibodies were used at a dilution of 1:50.

### ELISA

Basal prostaglandin E2 levels in culture supernatants from PDEs were quantified using a PGE_2_ ELISA kit (Cayman Chemical, 514010) according to the manufacturer’s instructions. Supernatants were collected after 24 hrs of culture, and PGE_2_ concentrations were determined by enzyme linked immunosorbent assay following the provided protocol.

### Cytokine profiling

After 24 hrs of *ex vivo* coculture of PDEs, culture supernatants were collected, centrifuged at 400 g for 10 min at 4 °C to remove residual cells and debris, and stored at -80 °C until analysis. Soluble mediators were quantified using BD Cytometric Bead Array assays (BD Biosciences) following the manufacturer’s instructions. Analytes included granzyme A (BD Biosciences Cat# 560299, RRID:AB_2869330), granzyme B (BD Biosciences Cat# 560304, RRID:AB_2869331), IFN γ (BD Biosciences Cat# 558269, RRID:AB_2869127), TNFα (BD Biosciences Cat# 560112, RRID:AB_2869306), CXCL9 (BD Biosciences Cat# 558341, RRID:AB_2869166), CXCL10 (BD Biosciences Cat# 558280, RRID:AB_2869136), CCL5 (BD Biosciences Cat# 558324, RRID:AB_2869154), G-CSF (BD Biosciences Cat# 558326, RRID:AB_2869156), GM-CSF (BD Biosciences Cat# 558335, RRID:AB_2869163), VEGF (BD Biosciences Cat# 558336, RRID:AB_2869164), IL-2 (BD Biosciences Cat# 558270, RRID:AB_2869128), IL-4 (BD Biosciences Cat# 558272, RRID:AB_2869129), IL-6 (BD Biosciences Cat# 558276, RRID:AB_2869132), IL-10 (BD Biosciences Cat# 558274, RRID:AB_2869131), and IL-1β (BD Biosciences Cat# 558279, RRID:AB_2869135) using the corresponding BD CBA kits. For each condition, 17 µL of supernatant was processed per assay according to the recommended protocol. Samples were acquired on a BD LSR Fortessa flow cytometer, and data were analyzed using FlowJo software version 10.9.0 (FlowJo LLC).

### Immunohistochemistry

Immunofluorescence analyses were performed on spheroids, PDTs, and PDEs before coculture to assess MSLN antigen expression and after coculture to evaluate treatment induced DNA damage (Sigma Aldrich, 11684809910) and apoptosis (cleaved-caspase 3). Following short term coculture, samples were fixed in 4% formalin at 4 °C overnight. Fixed tissues were embedded in HistoGel (Thermo Fisher Scientific)and processed using standard dehydration and clearing procedures prior to paraffin embedding.

Paraffin embedded blocks were sectioned at 4 µm thickness, with sections cut adjacent to the corresponding hematoxylin and eosin stained slide. Immunofluorescence staining was performed following the laboratory standard workflow. Sections were deparaffinized and antigen retrieval was carried out in Tris EDTA buffer at pH 9.0. Sections were permeabilized and blocked in PBS 2% BSA 0.03% Triton-X 100 for 1 h at room temperature, then incubated overnight at 4 °C with primary antibodies diluted PBS 2% BSA. Primary antibodies included mouse anti-panCK and rabbit anti-mesothelin (R&D systems, MAB32651), both used at a 1:100 dilution. Alexa Fluor conjugated secondary antibodies were then incubated for 1 h at room temperature. Nuclei were counterstained and sections were mounted using a DAPI containing mounting medium. Whole slide images were acquired using a Zeiss Axioscan 7 scanner at 20× magnification. Quantification of cytokeratin positive areas, mesothelin expression, and marker colocalization was performed using QuPath (RRID:SCR_018257) software v0.5.0 with identical analysis parameters applied across all sections.

### TUNEL assay

Tunel staining was performed on paraffin embedded sections using the In Situ Cell Death Detection Kit according to the manufacturer’s instructions. Briefly, antigen retrieval was carried out by incubation with proteinase K at 20 µg/mL for 15 min at 37 °C. Positive control sections were treated with DNase I at 3 U/mL (New England Biolabs, M0303) for 10 min at room temperature. Tissue sections were subsequently incubated with TUNEL reaction mixture for 1 h at 37 °C in a humidified chamber. After staining, sections were mounted using a DAPI containing mounting medium (Invitrogen, P36931). Whole slide fluorescence images were acquired using a Zeiss Axioscan 7 slide scanner at 20× magnification. Quantification of TUNEL positivity was performed using QuPath software version 0.5.0 by calculating the proportion of TUNEL positive nuclei relative to total DAPI positive nuclei across the entire explant area. For each patient, control and treated conditions were compared using a paired t-test in GraphPad Prism 8, with three to four explants per treatment condition and multiple regions of interest analyzed per tumor.

### Volumetric immunofluorescence microscopy imaging

To quantify exogenous T cell infiltration, MSLN-BiTE T cells and UT T cells were pre-labeled with CellTrace Yellow prior to coculture following manufacturer’s instructions. Tumor samples were cocultured with labeled T cells for 24 hrs, after which tissues were fixed in 4% formalin solution overnight at 4 °C. Following fixation, samples were permeabilized in PBS 0.3% Triton-X 100 for 24 hrs at room temperature under gentle agitation. Nonspecific binding sites were blocked using PBS 3% BSA 0.3% Triton-X 100 (blocking buffer) for 6 hrs at room temperature. Tissues were subsequently incubated with primary antibodies diluted in blocking buffer, including anti-panCK (1:200) and anti CD8 (1:150).

Volumetric imaging was performed using two-photon laser scanning microscopy. Tiled three dimensional image stacks were acquired to cover the entire tumor area in the XY plane, with a Z step size of 15 µm, generating volumetric datasets spanning a depth of approximately 500 to 800 µm. All image stacks were imported into Imaris software v10.0 (RRID:SCR_007370) and analyzed using surface detection and surface creation tools to generate three dimensional reconstructions of regions of interest and to quantify exogenous T cell infiltration within tumor compartments.

### *Ex vivo* characterization of patient response

*Ex vivo* functional stratification was performed at the patient level using tumor reactive T cell activation as the primary endpoint. Following PDE coculture, CD8^+^ T cell activation was quantified by flow cytometry as the frequency of CD8^+^CD137^+^ events within viable CD45^+^CD3^+^ T cells. For each patient, an activation fold change (FC) was computed by normalizing the MSLN-BiTE condition to the UT T cell control.

Patients were classified as responders (Rs) if FC≥ 2. Patients with FC <2 were classified as non-responders (NRs), irrespective of changes in soluble mediators. Cytokines and cytotoxic effector molecules quantified in culture supernatants were used as orthogonal functional readouts to describe response programs but were not used to assign responder status.

### Spatial transcriptome analysis: FFPE CytAssist Visium

For each FFPE block, one section of 5um-thick was freshly cut and placed onto a Superfrost+ adhesion slide (Epredia). Slides were then heated at 42 °C for 3 h on a thermocycler using the Low-Profile Thermocycler Adaptor (10XGenomics). Slides were stored at 4°C up to 2 weeks if not processed immediately. The day of experiment, slides were deparaffinized, H&E stained, imaged and decrosslinked following the 10X Genomics protocol “Visium CytAssist for FFPE” (CG000520). Image acquisition was done with the slide scanner Axioscan 7 (Zeiss). When slides were ready for CytAssist transfer, tissues were processed according to the protocol “Visium CytAssist Spatial Gene Expression” (CG000495) and probe-based libraries were obtained. Quality Control and DNA quantification of final libraries were performed with the Fragment Analyzer (Agilent Technologies) and the Qubit High-Sensitivity dsDNA assay kit (InVitrogen), respectively. Libraries were sequenced on AVITI (Element Biosciences) following 10X parameters recommendation (Paired-end, dual indexing: Read 1: 28 cycles; i7 Index: 10 cycles; i5 Index: 10cycles; Read 2: 50 cycles).

### Spatial transcriptomics data processing

Raw sequencing data were processed using Space Ranger v3.1.3 (10x Genomics) with default parameters for alignment to the GRCh38 reference genome and gene quantification. Processed count matrices and spatial coordinates were imported into R (v4.3) using the Load10X_Spatial function from Seurat (v5.0). Quality control was performed by manually excluding off-target spots located outside the H&E staining area. For each spot, mitochondrial and ribosomal gene content was calculated using the PercentageFeatureSet function with pattern matching for genes prefixed with MT- and RP[SL], respectively. Data normalization was performed using SCTransform with variance stabilization. Dimensionality reduction was conducted via principal component analysis (PCA) retaining 30 components based on elbow plot assessment. Neighborhood graphs were constructed using FindNeighbors, and unsupervised clustering was performed across multiple resolutions (0.1-10) using FindClusters. Non-linear dimensionality reduction was performed using both t-distributed stochastic neighbor embedding (t-SNE) and uniform manifold approximation and projection (UMAP) with a minimum distance parameter of 0.75. Spatially variable features were identified using Moran’s I statistic computed on the top 1,000 variable features. Integration features (n = 5,000) were selected using SelectIntegrationFeatures for downstream analyses.

### Cell type deconvolution

Cell type composition of each Visium spot was estimated using robust cell type decomposition (RCTD) from the spacexr package. A single-cell RNA sequencing reference atlas was constructed from high-grade serous ovarian cancer samples as described in Vázquez-García et al. (Nature, 2022). Cell type annotations were refined using the nomenclature and classification strategy established in Barras et al. (Science Immunology, 2024). The reference dataset was SCTransform-normalized prior to deconvolution. RCTD was executed in full mode (doublet_mode = “full”) allowing for multi-cell type assignment per spot, with analyses performed independently for each sample to account for inter-sample variability. Deconvolution was conducted at two levels of granularity: main cell lineages and fine-grained cell type annotations. Cell type proportion estimates were stored as separate assays within Seurat objects, and the dominant cell type for each spot was assigned based on maximum proportion values. Results were visualized using SpatialDimPlot.

### Tissue segmentation

Spatial spots were classified into three tissue microenvironment categories (Tumor, Stroma, Mixed) based on RCTD-derived cell type proportions. Spots were designated as tumor regions if malignant cell proportion exceeded 20%. Stromal regions were identified when CAF proportion exceeded 20% or when the combined stromal fraction (CAFs, endothelial cells, and pericytes) exceeded 30%. Spots meeting both tumor and stromal criteria were classified as mixed regions. To ensure spatial coherence of annotations, a neighborhood-based refinement algorithm was applied using the Giotto hexagonal spatial network. Isolated spots (i.e., those classified differently from all neighboring spots) were reclassified as mixed regions when surrounded entirely by spots of an opposing category, requiring a minimum of two neighbors for reclassification. This approach minimized spurious single-spot classifications while preserving biologically meaningful boundaries between tissue compartments.

### Pathway activity scoring

Pathway activity scores were computed using Gene Set Variation Analysis (GSVA) with a Gaussian kernel cumulative density function appropriate for SCTransform-normalized continuous expression data. Log-normalized expression matrices from the Spatial assay were used as input. A comprehensive collection of 2,420 gene sets was interrogated, derived from the following sources: MSigDB Hallmark gene sets, Reactome pathways, KEGG pathways, BioCarta pathways, immune cell type-specific signatures derived from single-cell studies and extracted from Lowery et al. (Science. 2022) and in-house curated signatures. Minimum and maximum gene set sizes were set to 5 and 500 genes, respectively. Signature scores were stored as a separate assay in Seurat objects for downstream comparative analyses.

### Statistical analysis of pathway scores

Comparative analyses of pathway activity scores were performed across experimental conditions (Untreated, MSLN-BiTE, MSLN-BiTE + COXi), tissue regions (Tumor, Stroma, Mixed), and clinical response categories (Rs, NRs). For each combination of variables, pathway scores were visualized using boxplots with individual spot values overlaid as jittered points. Statistical comparisons between treatment conditions were performed using pairwise Wilcoxon rank-sum tests with Benjamini-Hochberg correction for multiple testing. Significant differences were annotated using standard significance notation (*p < 0.05; **p < 0.01; ***p < 0.001). Median pathway scores were computed for each condition and summarized in heatmaps with column-wise z-score scaling. Hierarchical clustering was applied to columns (pathways) using Euclidean distance and complete linkage, while rows (condition combinations) were ordered by tissue region, response status, and treatment to facilitate biological interpretation.

### Differential gene expression analysis

Differential gene expression (DGE) analysis comparing MSLN-BiTE + COXi versus untreated conditions was performed using a dual-level analytical framework integrating spot-level and sample-level pseudobulk approaches, stratified by tissue region and clinical response. For spot-level analysis, raw counts from the Spatial assay were used. A linear model was fitted using limma-voom with the design matrix including patient as a blocking factor to account for paired samples (∼Patient + Treatment). Normalization factors were calculated using the trimmed mean of M-values (TMM) method, followed by voom transformation to estimate observation weights. Empirical Bayes moderation was applied to obtain moderated t-statistics and p-values. For pseudobulk analysis, counts were aggregated across all spots within each sample-treatment combination by summing raw counts. The same linear modeling approach was applied to pseudobulk data with patient as a covariate. P-values were adjusted using the Benjamini-Hochberg method for spot-level data and nominal p-values were used for pseudobulk analysis given the limited sample size.

### Pathway enrichment analysis of differentially expressed genes

Gene set enrichment analysis was performed on each gene significance category using the clusterProfiler enricher function. The same comprehensive gene set collection used for GSVA was employed as the background universe. Enrichment p-values were adjusted using the Benjamini-Hochberg method. For each significance category, the top enriched pathways were identified. Results were summarized in heatmaps displaying -log10(adjusted p-value) for the most significantly enriched pathways across all categories. Columns were hierarchically clustered while rows were ordered by significance category to facilitate interpretation of pathway associations with specific expression patterns. Separate analyses were conducted for responders and non-responders to identify response-specific transcriptional programs induced by treatment.

### Data visualization

Spatial feature plots were generated using SpatialFeaturePlot and SpatialDimPlot functions from Seurat. Heatmaps were produced using the pheatmap package with column-wise z-score scaling where indicated. Boxplots and scatter plots were created using ggplot2 with statistical annotations added via ggpubr. For enrichment analyses, treeplot visualizations of term similarities were generated using the enrichplot package when sufficient pathway terms were identified. All analyses were performed in R (v4.3) with visualization using ggplot2 (v3.4) and pheatmap (v1.0.12).

### Software and packages

The following software and R packages were used for spatial transcriptomics analysis: Space Ranger (10x Genomics) for raw data processing; Seurat (v5.0) for data preprocessing, normalization, and visualization; spacexr for RCTD deconvolution; Giotto for spatial network analysis; GSVA for pathway activity scoring; limma for differential expression analysis; clusterProfiler and enrichplot for pathway enrichment analysis; rstatix for statistical testing; pheatmap for heatmap visualization; and ggplot2 with ggpubr for data visualization and statistical annotation.

## References

1 Rosenberg SA, Restifo NP. Adoptive cell transfer as personalized immunotherapy for human cancer. Science 2015; 348 (6230): 62–68.

2 Pearce OMT, Delaine-Smith RM, Maniati E et al. Deconstruction of a Metastatic Tumor Microenvironment Reveals a Common Matrix Response in Human Cancers. Cancer Discov 2018; 8 (3): 304–319.

3 Ferri-Borgogno S, Zhu Y, Sheng J et al. Spatial Transcriptomics Depict Ligand-Receptor Cross-talk Heterogeneity at the Tumor-Stroma Interface in Long-Term Ovarian Cancer Survivors. Cancer Res 2023; 83 (9): 1503–1516.

4 Chap BS, Rayroux N, Grimm AJ et al. Crosstalk of T cells within the ovarian cancer microenvironment. Trends Cancer 2024; 10 (12): 1116–1130.

5 Voabil P, de Bruijn M, Roelofsen LM et al. An ex vivo tumor fragment platform to dissect response to PD-1 blockade in cancer. Nat Med 2021; 27 (7): 1250–1261.

6 Wu KZ, Adine C, Mitriashkin A et al. Making In Vitro Tumor Models Whole Again. Adv Healthc Mater 2023; 12 (14): e2202279.

7 Lopez E, Kamboj S, Chen C et al. In Vitro Models of Ovarian Cancer: Bridging the Gap between Pathophysiology and Mechanistic Models. Biomolecules 2023; 13 (1).

8 Mui M, Clark M, Vu T et al. Use of patient-derived explants as a preclinical model for precision medicine in colorectal cancer: A scoping review. Langenbecks Arch Surg 2023; 408 (1): 392.

9 Templeton AR, Jeffery PL, Thomas PB et al. Patient-Derived Explants as a Precision Medicine Patient-Proximal Testing Platform Informing Cancer Management. Front Oncol 2021; 11: 767697.

10 Ma L, Bai M, Wang Y et al. COX-2 inhibition synergizes with radioimmunotherapy by promoting TCF1(+)CD8(+) T cell infiltration in NSCLC. Cancer Biol Med 2025; 22 (12): 1537–1543.

11 Ye Y, Wang X, Jeschke U et al. COX-2-PGE(2)-EPs in gynecological cancers. Arch Gynecol Obstet 2020; 301 (6): 1365–1375.

12 Ghisoni E, Benedetti F, Minasyan A et al. Myeloid cell networks govern re-establishment of original immune landscapes in recurrent ovarian cancer. Cancer Cell 2025; 43 (8): 1568–1586 e1510.

13 Banerjee S, Ghisoni E, Wolfer A, et al. Bevacizumab, Atezolizumab, and Acetylsalicylic Acid in Recurrent, Platinum-Resistant Ovarian Cancer: The EORTC 1508-GCG Phase II Study. Clin Cancer Res 2025; 31 (11): 2145–2153.

14 Elewaut A, Estivill G, Bayerl F et al. Cancer cells impair monocyte-mediated T cell stimulation to evade immunity. Nature 2025; 637 (8046): 716–725.

15 Zelenay S, van der Veen AG, Bottcher JP et al. Cyclooxygenase-Dependent Tumor Growth through Evasion of Immunity. Cell 2015; 162 (6): 1257–1270.

16 Lacher SB, Dorr J, de Almeida GP et al. PGE(2) limits effector expansion of tumour-infiltrating stem-like CD8(+) T cells. Nature 2024; 629 (8011): 417–425.

17 Morotti M, Grimm AJ, Hope HC et al. PGE(2) inhibits TIL expansion by disrupting IL-2 signalling and mitochondrial function. Nature 2024; 629 (8011): 426–434.

18 Pelly VS, Moeini A, Roelofsen LM et al. Anti-Inflammatory Drugs Remodel the Tumor Immune Environment to Enhance Immune Checkpoint Blockade Efficacy. Cancer Discov 2021; 11 (10): 2602–2619.

19 Kosti P, Abram-Saliba J, Pericou-Troquier L et al. Potent and durable control of mesothelin-expressing tumors by a novel T cell-secreted bi-specific engager. J Immunother Cancer 2025; 13 (3).

20 Katt ME, Placone AL, Wong AD et al. In Vitro Tumor Models: Advantages, Disadvantages, Variables, and Selecting the Right Platform. Front Bioeng Biotechnol 2016; 4: 12.

21 Ye Q, Song DG, Poussin M et al. CD137 accurately identifies and enriches for naturally occurring tumor-reactive T cells in tumor. Clin Cancer Res 2014; 20 (1): 44–55.

22 Kim M, Mun H, Sung CO et al. Patient-derived lung cancer organoids as in vitro cancer models for therapeutic screening. Nat Commun 2019; 10 (1): 3991.

23 Nanki Y, Chiyoda T, Hirasawa A et al. Patient-derived ovarian cancer organoids capture the genomic profiles of primary tumours applicable for drug sensitivity and resistance testing. Sci Rep 2020; 10 (1): 12581.

24 Vias M, Morrill Gavarro L, Sauer CM et al. High-grade serous ovarian carcinoma organoids as models of chromosomal instability. Elife 2023; 12.

25 Thorel L, Perreard M, Florent R et al. Patient-derived tumor organoids: a new avenue for preclinical research and precision medicine in oncology. Exp Mol Med 2024; 56 (7): 1531–1551.

26 Barras D, Ghisoni E, Chiffelle J et al. Response to tumor-infiltrating lymphocyte adoptive therapy is associated with preexisting CD8(+) T-myeloid cell networks in melanoma. Sci Immunol 2024; 9 (92): eadg7995.

27 Duraiswamy J, Turrini R, Minasyan A et al. Myeloid antigen-presenting cell niches sustain antitumor T cells and license PD-1 blockade via CD28 costimulation. Cancer Cell 2021; 39 (12): 1623–1642 e1620.

28 Ibanez-Molero S, Veldman J, Simon Nieto J et al. Tumour-reactive heterotypic CD8 T cell clusters from clinical samples. Nature 2026; 649 (8096): 467–476.

29 Thomas A, Chen Y, Steinberg SM et al. High mesothelin expression in advanced lung adenocarcinoma is associated with KRAS mutations and a poor prognosis. Oncotarget 2015; 6 (13): 11694–11703.

30 Magalhaes I, Fernebro J, Abd Own S et al. Mesothelin Expression in Patients with High-Grade Serous Ovarian Cancer Does Not Predict Clinical Outcome But Correlates with CD11c(+) Expression in Tumor. Adv Ther 2020; 37 (12): 5023–5031.

31 Weidemann S, Gagelmann P, Gorbokon N et al. Mesothelin Expression in Human Tumors: A Tissue Microarray Study on 12,679 Tumors. Biomedicines 2021; 9 (4).

32 Chiffelle J, Barras D, Petremand R et al. Tumor-reactive T cell clonotype dynamics underlying clinical response to TIL therapy in melanoma. Immunity 2024; 57 (10): 2466-2482 e2412.

33. Chiello JL, Shaikh N, Jacobi J et al. BiTE-secreting T cells rationally combine with PD-1 blockade and vaccine boosting to reshape antitumor immunity in ovarian cancer. Mol Ther 2026; 34 (1): 455–478.

34 T Kim E, Kim JH, Park EY et al. The Efficacy and Safety of Folate Receptor alpha-Targeted Antibody-Drug Conjugate Therapy in Patients With High-Grade Epithelial Ovarian, Primary Peritoneal, or Fallopian Tube Cancers: A Systematic Review and Meta-Analysis. Cancer Med 2024; 13 (21): e70392.

35 McDermott MSJ, O’Brien NA, Hoffstrom B et al. Preclinical Efficacy of the Antibody-Drug Conjugate CLDN6-23-ADC for the Treatment of CLDN6-Positive Solid Tumors. Clin Cancer Res 2023; 29 (11): 2131–2143.

36 Gaspar M, Natoli M, Castan L et al. An affinity-modulated T cell engager targeting Claudin 18.2 shows potent anti-tumor activity with limited cytokine release. J Immunother Cancer 2025; 13 (8).

37 McGray AJR, Chiello JL, Tsuji T et al. BiTE secretion by adoptively transferred stem-like T cells improves FRalpha+ ovarian cancer control. J Immunother Cancer 2023; 11 (6).

