## Supplemental Figures for "Actionable spatial prostanoid barriers constrain BiTE-driven adoptive T cell immunity in intact human tumors"

Supplementary figure 1

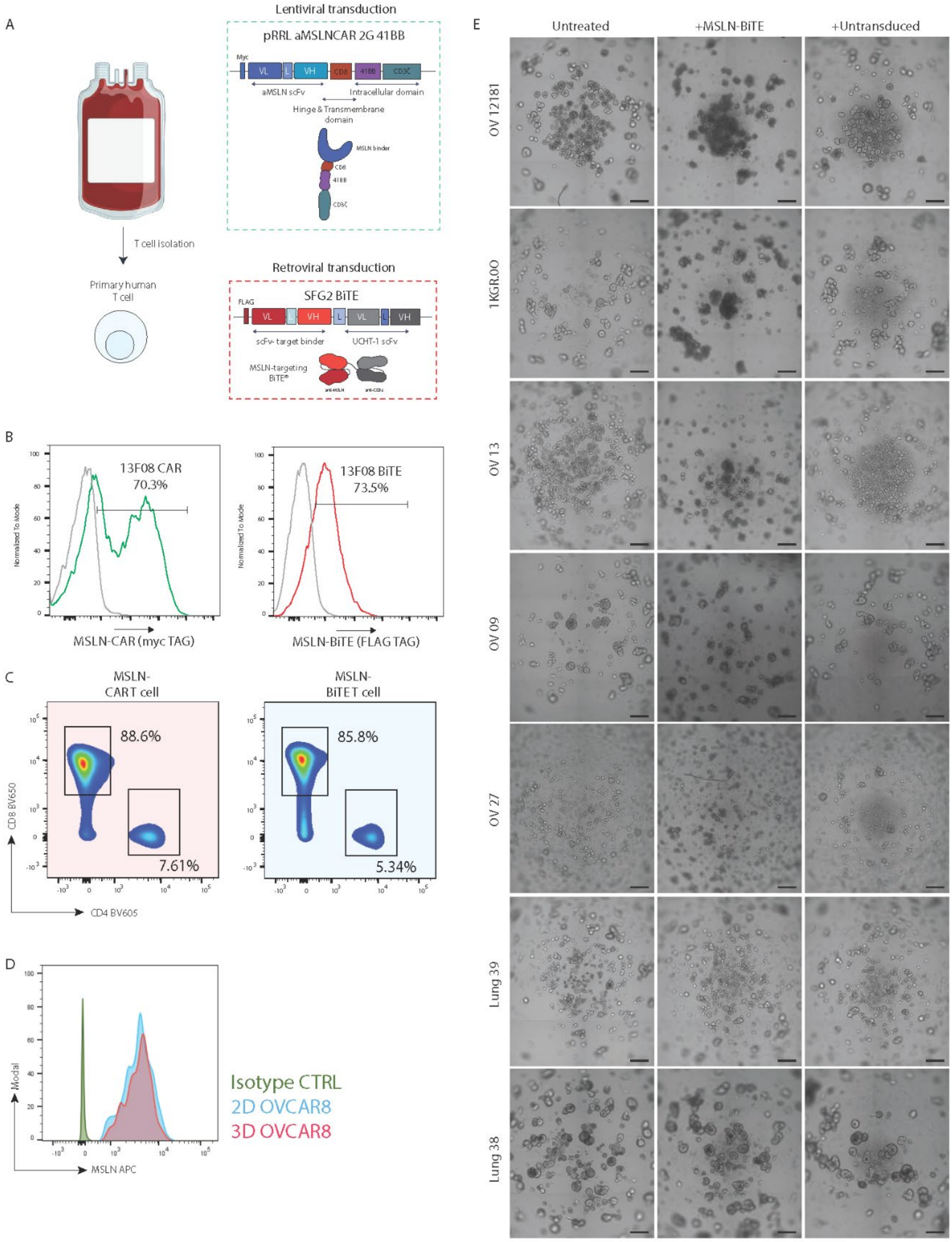

**Supplementary figure 1:**

**Generation and validation of MSLN-directed CAR T cells and BiTE-secreting T cells for antigen-defined 3D assays. A,** Schematic workflow MSLN-redirected T cell products generation from primary human T cells. Cells were transduced with a lentiviral second-generation anti-MSLN CAR construct containing a 4-1BB costimulatory domain (aMSLN-CAR 2G 41BB; myc-tagged for detection) or with a retroviral vector encoding a secreted anti-MSLN x anti-CD3 BiTE (SFG2 BiTE; FLAG-tagged for detection). **B,** Representative flow cytometry histograms confirming expression of the 13F08 CAR (myc tag) and the 13F08 BiTE (FLAG tag). Percent positive cells are indicated on each histogram. **C,** Representative flow cytometry plots showing $CD4^{+}$ and $CD8^{+}$ composition of transduced MSLN-CAR T cells and MSLN-BiTE T cells, with gated frequencies indicated. **D,** Flow cytometry assessment of surface MSLN expression on OVCAR-8 cultured in 2D monolayer or as 3D spheroids, overlaid with the corresponding isotype control. **E,** Representative bright-field images from multiple PDTs cultures exposed to no treatment (Untreated), MSLN-BiTE, or UT T cells CTRL, illustrating inter-patient variability in baseline morphology and treatment-associated disruption. Scale bars = 250 μm. Figure generated with BioRender.

Supplementary figure 2

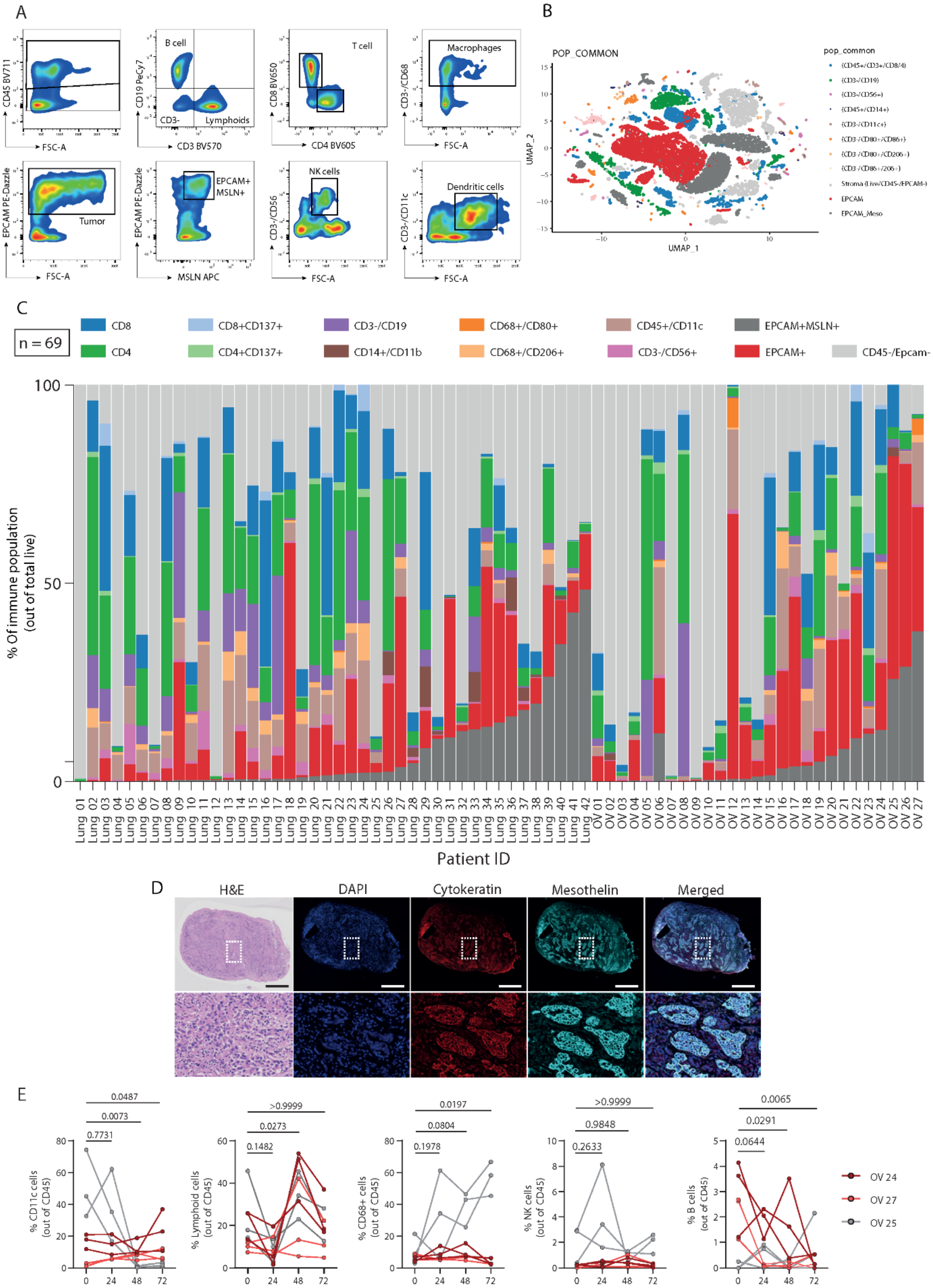

**Supplementary figure 2:**

**Baseline cellular composition and longitudinal immune stability profiling of NSCLC and HGSOC explants by multiparameter flow cytometry and IF. A,** Representative flow cytometry gating strategy used to profile major cellular compartments in dissociated tumor specimens. Gates define $CD45^+$ immune cells and subsets including B cells ($CD19^+$), T cells ($CD3^+$ with $CD4^+$ and $CD8^+$ delineation and CD137 activation readouts), NK cells ($CD3^-CD56^+$), macrophages ($CD68^+$ with polarization markers such as $CD206^+$ and $CD80^+$), and dendritic cells ($CD45^+/CD11c^+$). In parallel, non-immune compartments were quantified by EPCAM and CD45 to resolve epithelial tumor cells, including the $EPCAM^+MSLN^+$ subset. **B,** UMAP visualization of single-cell cytometry data showing the distribution of phenotypic clusters across the profiled samples. Colors indicate the annotated cell populations listed in the legend, highlighting separation of epithelial, stromal, lymphoid, and myeloid compartments. **C,** Cohort-wide composition of tumor specimens (n = 69) across NSCLC and HGSOC lesions. Stacked bar plots show the relative frequencies of key immune and tumor compartments per patient, including $CD8^+$ T cells, $CD8^+CD137^+$ activated T cells, $CD4^+$ T cells, $CD4^+CD137^+$ activated T cells, $CD19^+$ B cells, $CD14^+$ myeloid cells, $CD68^+$ macrophage subsets ($CD80^+$ & $CD206^+$), $CD11c^+$ dendritic cells, NK cells ($CD3^-CD56^+$), $EPCAM^+$ tumor cells, $EPCAM^+MSLN^+$ tumor cells, and $CD45^-$/$Epcam^-$ fractions. **D,** Representative histology and IF characterization of a PDE section illustrating preservation of epithelial architecture and target expression. Whole-section and zoomed views are shown for H&E, nuclei (DAPI), CK (red), MSLN (cyan), and merged channels. **E,** Longitudinal assessment of immune compartment stability during PDE culture. Line plots show the frequencies of $CD11c^+$ cells, total lymphoid cells, $CD8^+$ T cells, NK cells, and B cells (each expressed as a fraction of $CD45^+$ cells) across 0, 24, 48, and 72 hrs for three independent lesions/patients. Each line represents one explant replicate. Replicate explants were averaged per patient per timepoint; statistics were performed on patient-level means (n = 3 patients) using one-way repeated-measures ANOVA with Dunnett's multiple-comparisons test versus 0 h (GraphPad Prism v10.1.2).

Supplementary figure 3

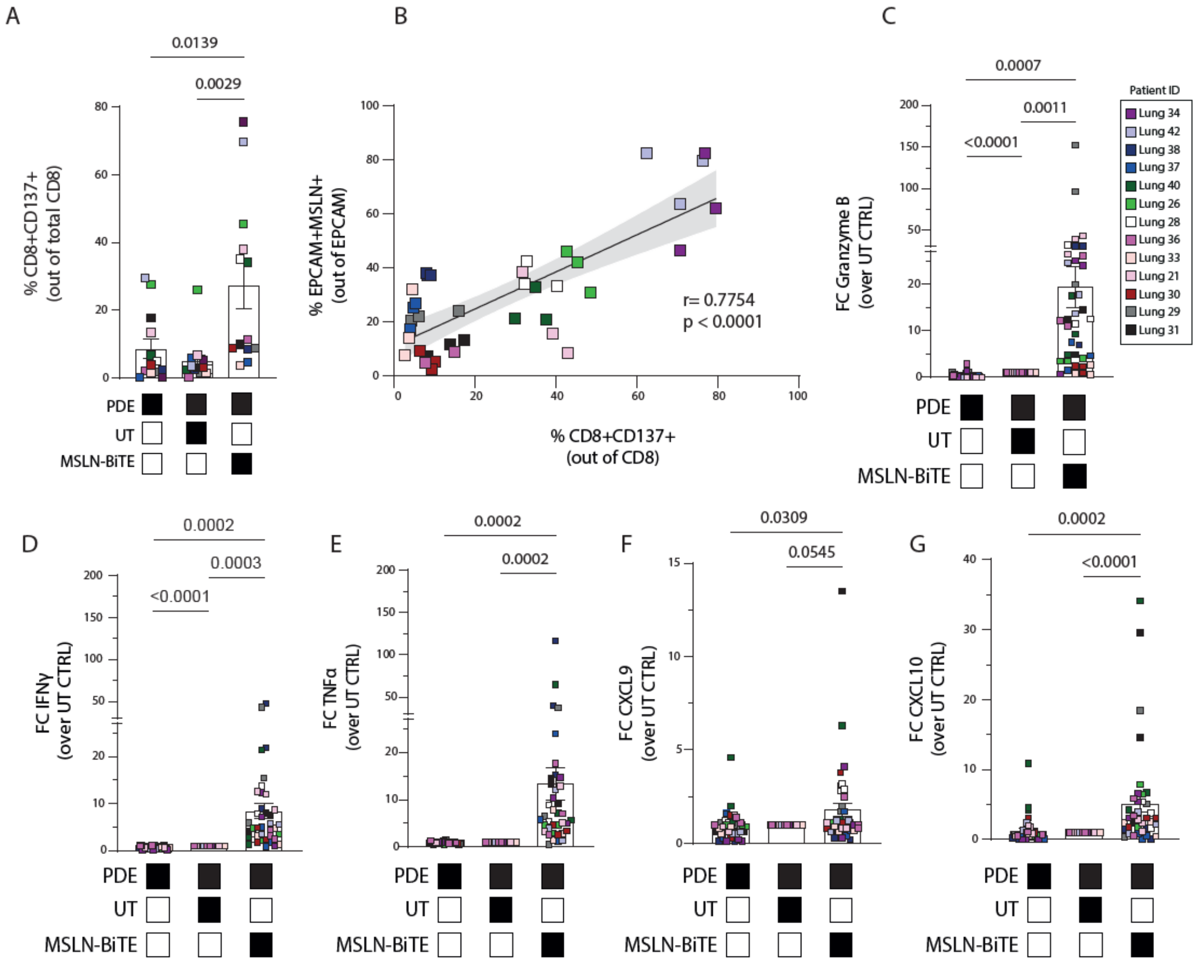

**Supplementary figure 3:**

**MSLN antigen abundance predicts MSLN-BiTE induced CD8 activation and effector mediator release in NSCLC explants. A,** Flow cytometry quantification of activated CD8 T cells, reported as the frequency of $CD8^+CD137^+$ cells among total $CD8^+$ T cells across untreated tissue and PDE cultures under the indicated conditions (PDE alone, PDE plus UT T cells, or PDE plus MSLN-BiTE T cells; condition matrix shown below). **B,** Pearson'r correlation plot describing association between target availability and functional activation. The percentage of $EPCAM^+MSLN^+$ cells among $EPCAM^+$ tumor cells is plotted against the frequency of $CD8^+CD137^+$ cells (out of $CD8^+$) across lesions; correlation coefficient (r) and P value are indicated. **C,** Granzyme B secretion measured in culture supernatants and expressed as fold change (FC) over the corresponding UT T cell control. IFNγ (**D**), TNFα (**E**), CXCL9 (**F**), and CXCL10 (**G**) quantified in supernatants and expressed as FC over UT CTRL across the same conditions. Each symbol represents one PDE fragment (technical replicate). For each lesion and condition, fragment values were averaged to generate a single lesion-level value. Statistical testing was performed on lesion-level values using one-way repeated-measures ANOVA with multiple-comparisons correction as indicated by the plotted contrasts (GraphPad Prism v10.1.2). Bars show mean ± SEM of lesion-level values.

Supplementary figure 4

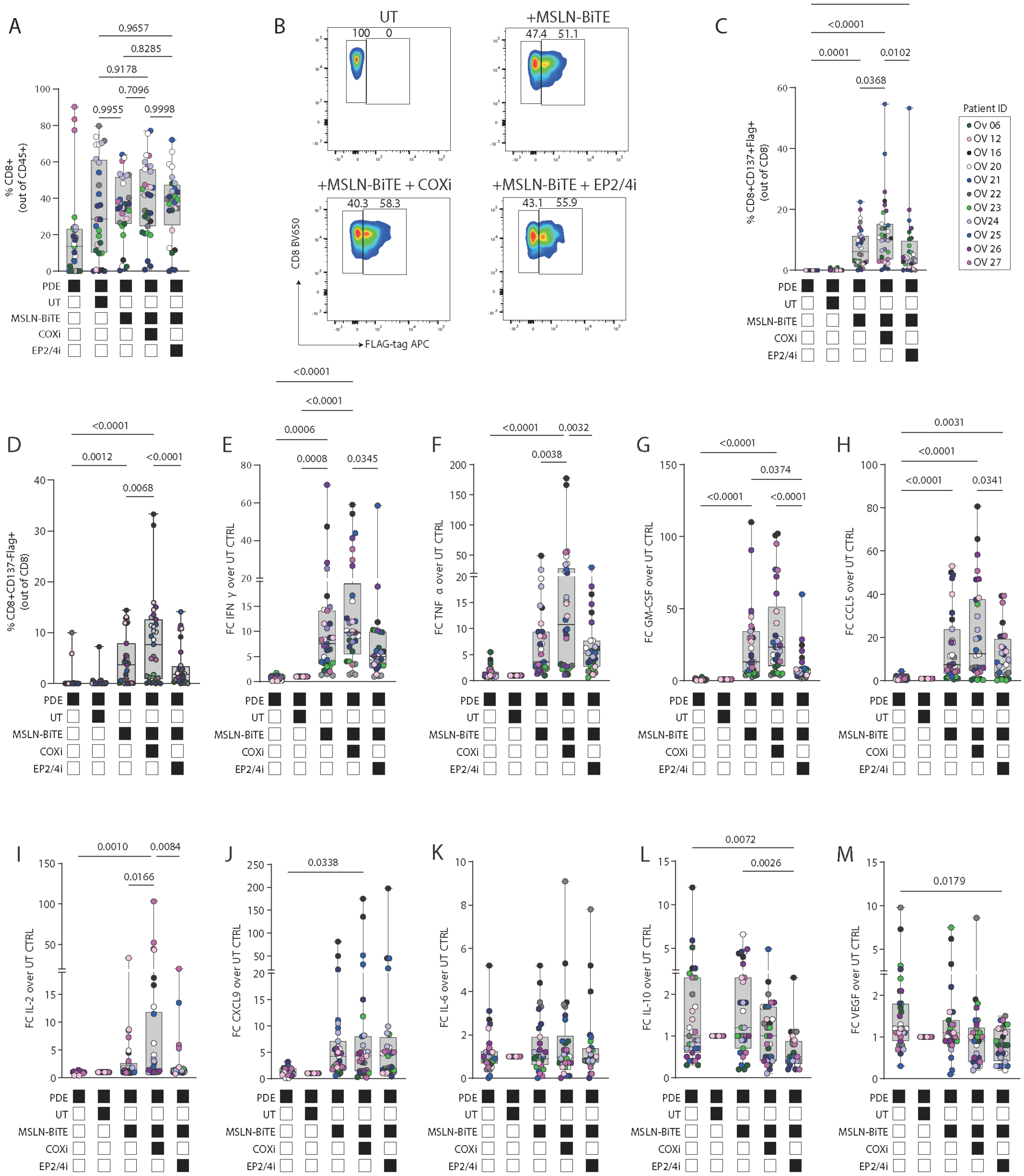

**Supplementary figure 4:**
**COX inhibition enhances lesion-level responses to MSLN-BiTE T cells in HGSOC explants.** HGSOC PDEs were co-cultured with UT or MSLN-BiTE-secreting T cells with or without COX inhibition (COXi) or EP2/EP4 antagonism (EP2/4i). Each dot represents one explant fragment (n = 2-3 per patient); boxplots show median and interquartile range and condition matrices indicate experimental groups. **A,** Total $CD8^+$ infiltration (%$CD8^+$ of $CD45^+$). **B,** Representative gating to distinguish exogenous $FLAG^+$ MSLN-BiTE T cells from $FLAG^-$ bystander/resident T cells. **C-D,** Frequencies of activated exogenous $CD8^+CD137^+FLAG^+$ cells (**C**) and activated bystander/resident $CD8^+CD137^+FLAG^-$ cells (**D**). **E-M,** Supernatant soluble factors expressed as fold-change over UT CTRL: IFNγ (**E**), TNFα (**F**), GM-CSF (**G**), CCL5 (**H**), IL-2 (**I**), CXCL9 (**J**), IL-6 (**K**), IL-10 (**L**), and VEGF (**M**). Each dot represents one patient fragment. Error bars indicate ± SEM. Statistics were performed using one-way repeated-measures ANOVA (GraphPad Prism v10.1.2); exact P values are shown.

Supplementary figure 5

A

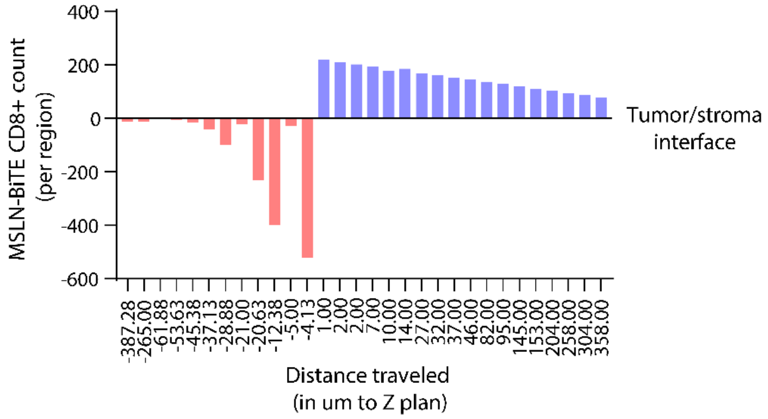


B

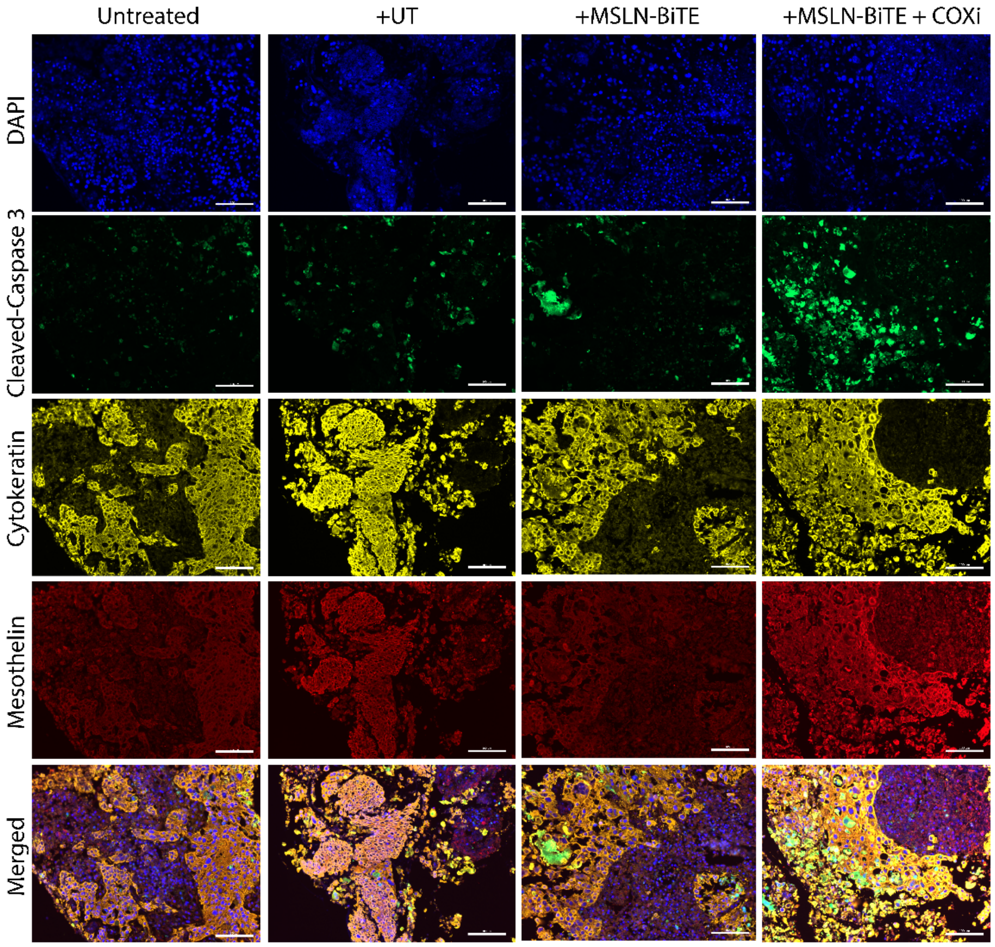

**Supplementary figure 5:**

**Depth-resolved distribution of MSLN-BiTE CD8 T cells across the tumor-stroma interface and tumor-restricted apoptosis. A,** Bar plot showing the depth profile of MSLN-BiTE $CD8^+$ T cell counts per segmented region as a function of distance along the z-axis, aligned to the tumor-stroma interface (0 µm). Values on either side of the interface reflect $CD8^+$ distribution across adjacent compartments. **B,** Representative IF images of PDE sections after 24 hrs co-culture under the indicated conditions (untreated, UT, MSLN-BiTE, MSLN-BiTE + COXi), stained for DAPI (blue), CC-3 (green), CK (yellow), MSLN (red), and merged channels, illustrating apoptosis concentrated within $CK^+MSLN^+$ tumor regions upon MSLN-BiTE treatment and further increased by COX inhibition; scale bar = 50 µm.

Supplementary figure 6

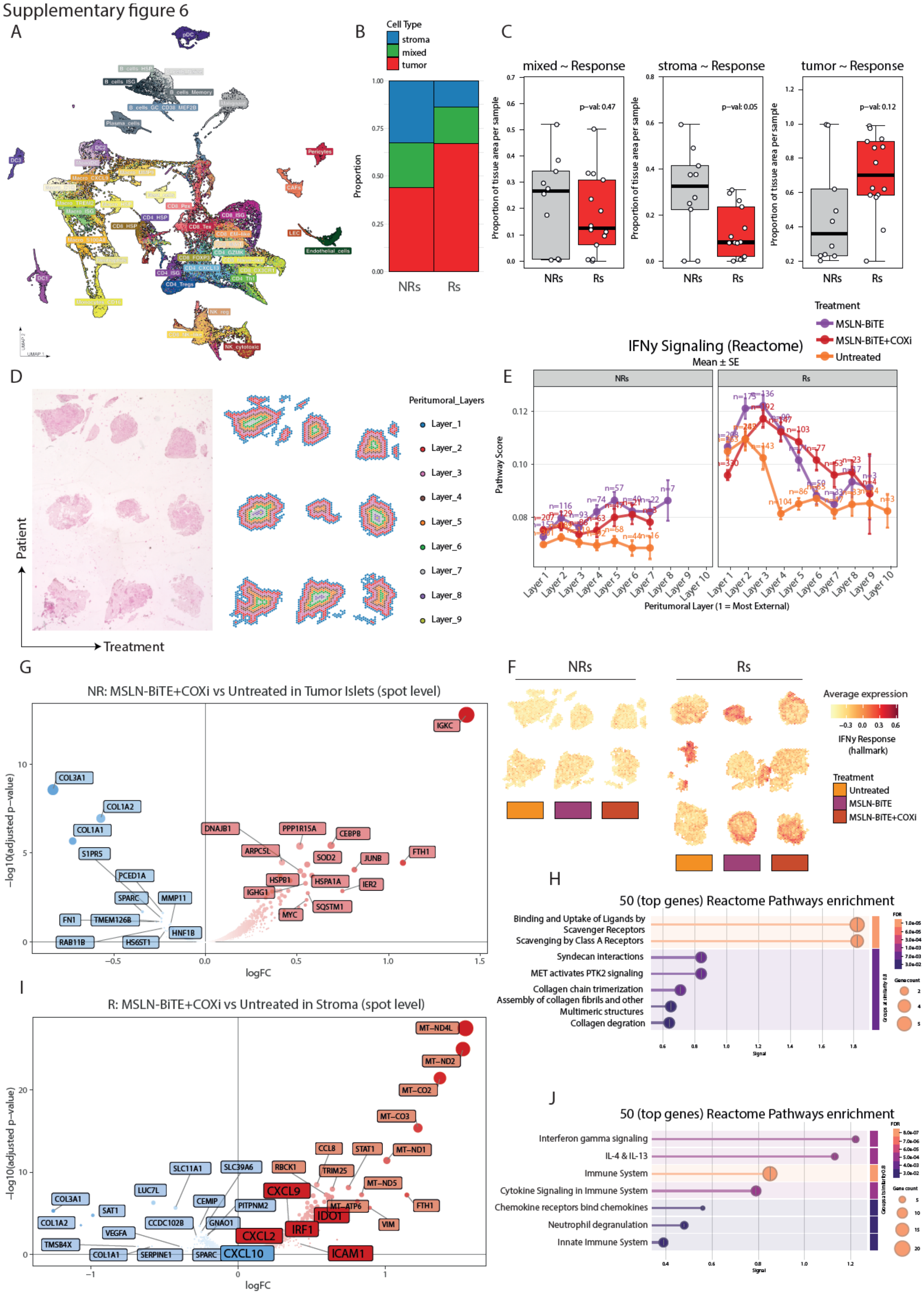

**Supplementary figure 6:**

**Spatial stratification of R and NR HGSOC PDEs reveals localized IFNγ programs and distinct compartmental remodeling. A,** UMAP of Visium spot deconvolution results showing the major inferred cell states across all samples, with clusters annotated by cell identity. **B,** Stacked bar plot comparing the relative proportions of tumor, mixed, and stromal tissue compartments in NR versus R lesions. **C,** Box plots showing the fraction of tissue classified as mixed, stroma, or tumor per sample in NR versus R lesions (p values indicated). **D,** Example of peritumoral spatial layering: H&E image (left) and corresponding concentric layer masks (right) defining successive spatial bands from the outer peritumoral region toward tumor islets. **E,** IFNγ signaling (Reactome) pathway scores plotted across peritumoral layers for NR (left) and R (right) lesions, stratified by treatment (Untreated, MSLN-BiTE, MSLN-BiTE + COXi). Mean ± SEM and spot numbers per layer are shown. **F,**) Spatial feature maps of an IFNγ response signature (Hallmark) across NR and R lesions under the indicated treatments, shown as average expression per spot (scale indicated). **G-H,** NR tumor-islet differential expression for MSLN-BiTE + COXi versus Untreated at spot level. **G,** Volcano plot with representative genes highlighted in NR tumor islet region. **H,** Reactome pathway enrichment of the top differentially expressed genes. **I-J,** Rs stromal differential expression for MSLN-BiTE + COXi versus Untreated at spot level. **I,** volcano plot with representative genes highlighted in responder stromal region. **J,** Reactome pathway enrichment of the top differentially expressed genes. Figure generated with BioRender.
